# Dorsal pulvinar inactivation leads to spatial selection bias without perceptual deficit

**DOI:** 10.1101/2022.12.30.520322

**Authors:** Kristin Kaduk, Melanie Wilke, Igor Kagan

## Abstract

The dorsal pulvinar has been implicated in visuospatial attentional and perceptual confidence processing. Pulvinar lesions in humans and monkeys lead to spatial neglect symptoms, including an overt spatial saccade bias during free choices. But it remains unclear whether disrupting the dorsal pulvinar during target selection that relies on a perceptual decision leads to a perceptual impairment or a more general spatial orienting and choice deficit. To address this question, we reversibly inactivated the unilateral dorsal pulvinar by injecting GABA-A agonist THIP while two macaque monkeys performed a color discrimination saccade task with varying perceptual difficulty. We used Signal Detection Theory and simulations to dissociate perceptual sensitivity (d-prime) and spatial selection bias (response criterion) effects. We expected a decrease in d-prime if dorsal pulvinar affects perceptual discrimination and a shift in response criterion if dorsal pulvinar is mainly involved in spatial orienting. After the inactivation, we observed response criterion shifts away from contralesional stimuli, especially when two competing stimuli in opposite hemifields were present. Notably, the d-prime and overall accuracy remained largely unaffected. Our results underline the critical contribution of the dorsal pulvinar to spatial orienting and action selection while showing it to be less important for visual perceptual discrimination.

## Introduction

Visual scenes contain multiple spatial locations that serve as potential saccades targets. Selecting a target in a complex scene requires efficiently perceiving and evaluating behaviorally relevant information at different spatial locations. Many studies investigating the neural processes of visuospatial target selection emphasize interactions in frontoparietal cortical networks (Corbetta and Shulman, 2011; Fiebelkorn et al., 2018; Adam et al., 2020). These direct cortical connections are paralleled by indirect routes through higher-order thalamic nuclei such as the pulvinar, raising the question of how the pulvinar contributes to the selection of behaviorally relevant stimuli to guide visuospatial decision-making (Halassa and Kastner, 2017; Sherman, 2017).

The primate pulvinar consists of several nuclei – anterior, medial, lateral and inferior - with distinct functional properties and connectivity profiles. The dorsal part of the pulvinar (dPul), the focus of the current study, encompasses the anterior and the medial pulvinar and the dorsal part of the lateral pulvinar (Gutierrez et al., 2000; Stepniewska, 2003; Arcaro et al., 2015; Baldwin and Bourne, 2017). These nuclei developed together with the association cortices in the course of primate evolution and are reciprocally connected to the parietal and prefrontal cortex, orbitofrontal cortex, insula, cingulate, superior and inferior temporal cortex (Kaas and Lyon, 2007; Preuss, 2007; Kaas and Baldwin, 2020). This extensive connectivity profile identifies the dorsal pulvinar as a unique brain hub well situated to interact with and modulate the circuitry involved in spatial attentional orienting and target selection (Grieve et al., 2000; Sherman and Guillery, 2002; Shipp, 2003; Saalmann and Kastner, 2015; Bourgeois et al., 2020; Kagan et al., 2021). Studies in human patients with unilateral thalamic lesions encompassing the pulvinar demonstrated deficits related to orienting or responding to perceptually or behaviorally salient stimuli in the contralesional hemifield (Danziger et al., 2001; Karnath et al., 2002; Arend et al., 2008b, 2008a; Lucas et al., 2019). Similar to neglect/extinction in patients, monkey studies using targeted reversible unilateral pharmacological inactivation of the dorsal pulvinar induced contralesional visuospatial deficits (Petersen et al., 1987; Desimone et al., 1990; Wilke et al., 2010, 2013; Komura et al., 2013). Such deficits manifest behaviorally in several ways. Firstly, the inactivation causes impairment of spatial attentional orienting to cued targets in contralesional hemifield, decreasing performance in detection or color-contingent manual response tasks (Petersen et al., 1987; Desimone et al., 1990). Secondly, the confidence about contralesional perceptual categorization but not the categorization itself has been reported to decrease after the inactivation, in the absence of competing distractors in the opposite hemifield (Komura et al., 2013). Thirdly, a target selection bias away from contralesional hemifield was observed in a free-choice saccade task (Wilke et al., 2010, 2013). Such inactivation-induced bias could be alleviated by presenting only a single target or increasing the reward for contralesional targets but less so by perceptual saliency manipulations (Wilke et al., 2013).

The above causal perturbation findings, and the results of electrophysiological recordings (Robinson and Petersen, 1992; Benevento and Port, 1995; Bender and Youakim, 2001; Dominguez-Vargas et al., 2017; Fiebelkorn et al., 2019; Schneider et al., 2019, 2023), on the one hand, implicate dPul in attentional allocation for perceptual processing, but on the other, are also compatible with a role in more general spatial orienting and selection bias. Different task demands might be one reason for such interpretational ambiguity. In particular, studies that used attentional cueing and perceptual discrimination employed paradigms where manual responses (e.g. button presses) were dissociated from the spatial position of the visual stimuli. At the same time, our previous choice tasks always required a saccade towards a peripheral target and did not involve difficult perceptual discrimination (Wilke et al., 2010, 2013; Dominguez-Vargas et al., 2017). Therefore, it remains unclear if perceptual factors contribute to contralesional visuospatial deficits under conditions of spatial competition and target-congruent saccade actions.

In the current study we used a color discrimination saccade selection task to address this question with two essential features. Firstly, we included easy and difficult (i.e. perceptually similar to a target) distractors that should not be selected with a saccade. Secondly, we introduced an option to maintain central fixation as a correct response when only distractor(s) were presented. The task involved three different stimulus types – single stimuli, double “same” stimuli (target-target or distractor-distractor) and double different stimuli. Single stimuli included a peripheral target or a distractor and a central fixation option, resulting in low spatial competition. The double stimuli included left and right peripheral stimuli and a fixation option, adding competition between hemifields. Using Signal Detection Theory (Stanislaw and Todorov, 1999; Macmillan and Creelman, 2004), we investigated whether unilateral dPul inactivation leads to a perceptual deficit or a spatial selection bias, similarly to recent cortical and superior colliculus studies (Luo and Maunsell, 2015, 2019; Lovejoy and Krauzlis, 2017; Crapse et al., 2018). Suppose the dorsal pulvinar is mainly involved in spatial orienting. In that case, we expected a shift in the criterion manifesting as a selection bias away from the contralesional hemifield, regardless of whether a target or a distractor is presented. But if the dorsal pulvinar is involved in discriminating targets from distractors, we expected a contralesional perceptual sensitivity deficit, manifesting as a decrease in d-prime. Furthermore, if dPul is mainly relevant for regulating the competition between hemifields, we expected a more substantial effect of inactivation in double stimuli conditions.

## Results

We investigated whether local unilateral injections of GABA-A agonist THIP suppressing the dorsal pulvinar neuronal activity (**Figure 1A**) cause a contralesional perceptual discrimination deficit or a spatial selection bias. Two monkeys performed a color discrimination task between red targets and distractors (orange stimuli as difficult distractors and yellow stimuli as easy distractors) in three stimulus type conditions (single stimuli, double same stimuli, double different stimuli, **Figure 1B**). We analyzed the following dependent variables: saccade latency, accuracy, d-prime (sensitivity), and response criterion (spatial choice bias). The main statistical analysis focused on the comparison between control vs. inactivation sessions.

**Figure 1.**
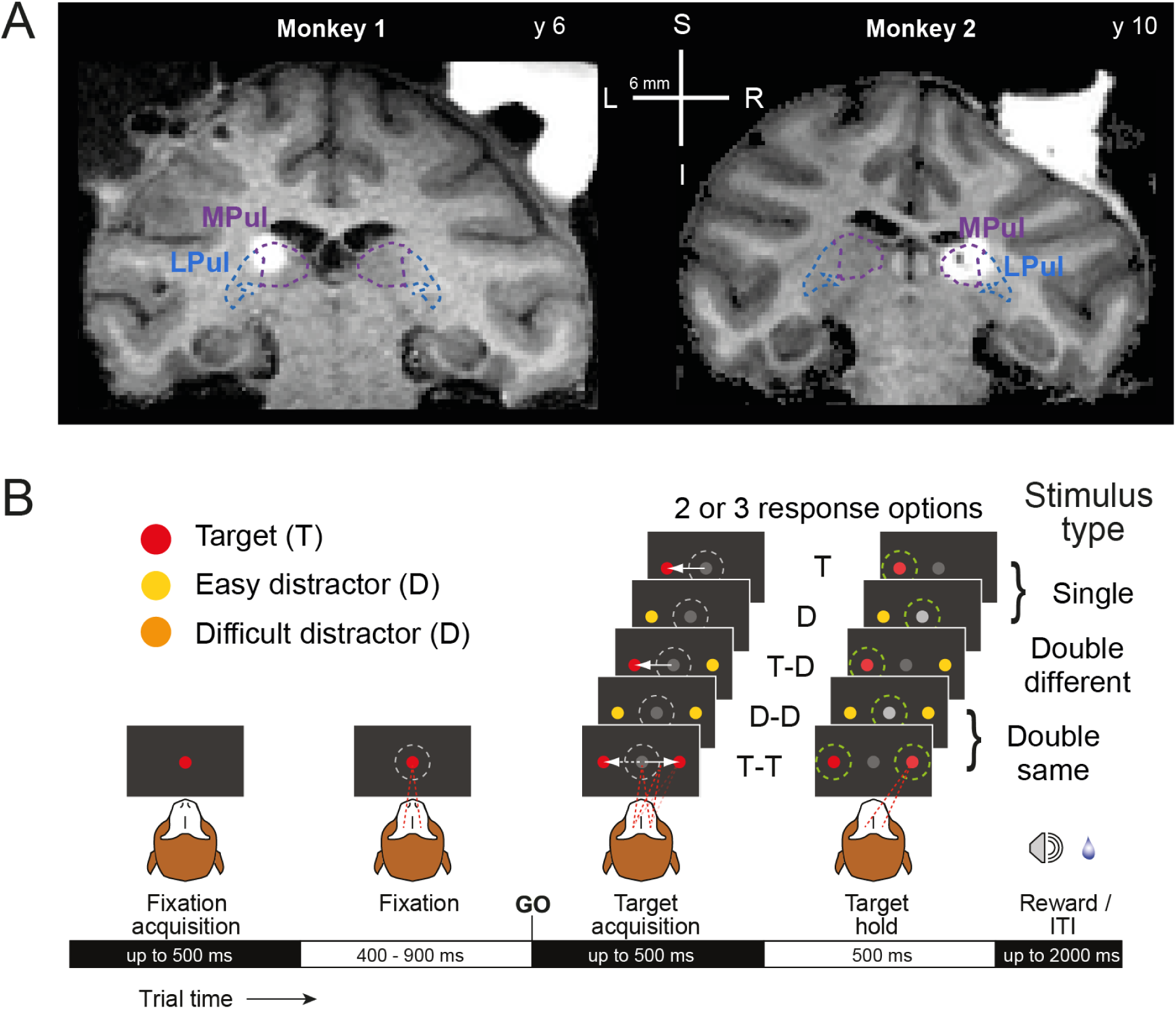
Inactivation sites and task design. (A) MR images show the inactivation sites visualized with co-injection of gadolinium MR contrast agent (ratio: 1:200 saline) ∼20 min following the injection (for M1: 2 μl and M2: 3 μl, respectively) and the overlaid borders of medial pulvinar (MPul) and lateral pulvinar (LPul). (B) Color discrimination saccade task where the perceptual difficulty was determined by the color similarity of the target (T, red) vs. distractor (D, easy - yellow, difficult - orange). Target or distractor was presented alone or with a second stimulus (distractor or target) in the opposite hemifield. Monkeys had to saccade to a target after the Go signal or continue fixating when only distractor(s) were presented. Green dashed circles in the target hold period denote correct and rewarded responses.

### The effect of dorsal pulvinar inactivation on saccade latency

The saccade latency is a sensitive measure of the effect of dorsal pulvinar inactivation. According to previous work, we expected either faster ipsilesional saccades (Wilke et al., 2010) or/and slower contralesional saccades (Wilke et al., 2013). A three-way mixed ANOVA (see details in **Suppl. Table S1**) was performed to compare the effect of the within-factors “Stimulus Type” (Single / Double Same / Double Different) and “Hemifield” (Contra- / Ipsilateral hemifield) and the between-factor “Perturbation” (Control / Inactivation sessions) on saccade latency. We found a statistically significant Stimulus type × Hemifield × Perturbation interaction for both monkeys (3-way interaction: M1: F(2,22) = 4.64, p = .021; M2: F(2,24) = 3.74, p = .039), as well as the Hemifield × Perturbation interaction for M1 (2-way interaction: M1: F(1,11) = 21.81, p = .001; M2: F(1,12) = 3.74, p = .08).

In agreement with our expectations about the inactivation effects, both monkeys significantly slowed down after dorsal pulvinar inactivation during contralesional target selection for both difficulty levels in all three stimulus types (independent t-test, **Table 1**, **Figure 2**). The consistent inactivation effect across stimulus types and difficulty levels on the contralesional saccade latency indicated a successful pharmacological manipulation. The saccade latency effects for ipsilesional target selection were less pronounced and only reached significance for the double same stimuli (independent t-test, **Table 1**, **Figure 2**).

**Figure 2.**
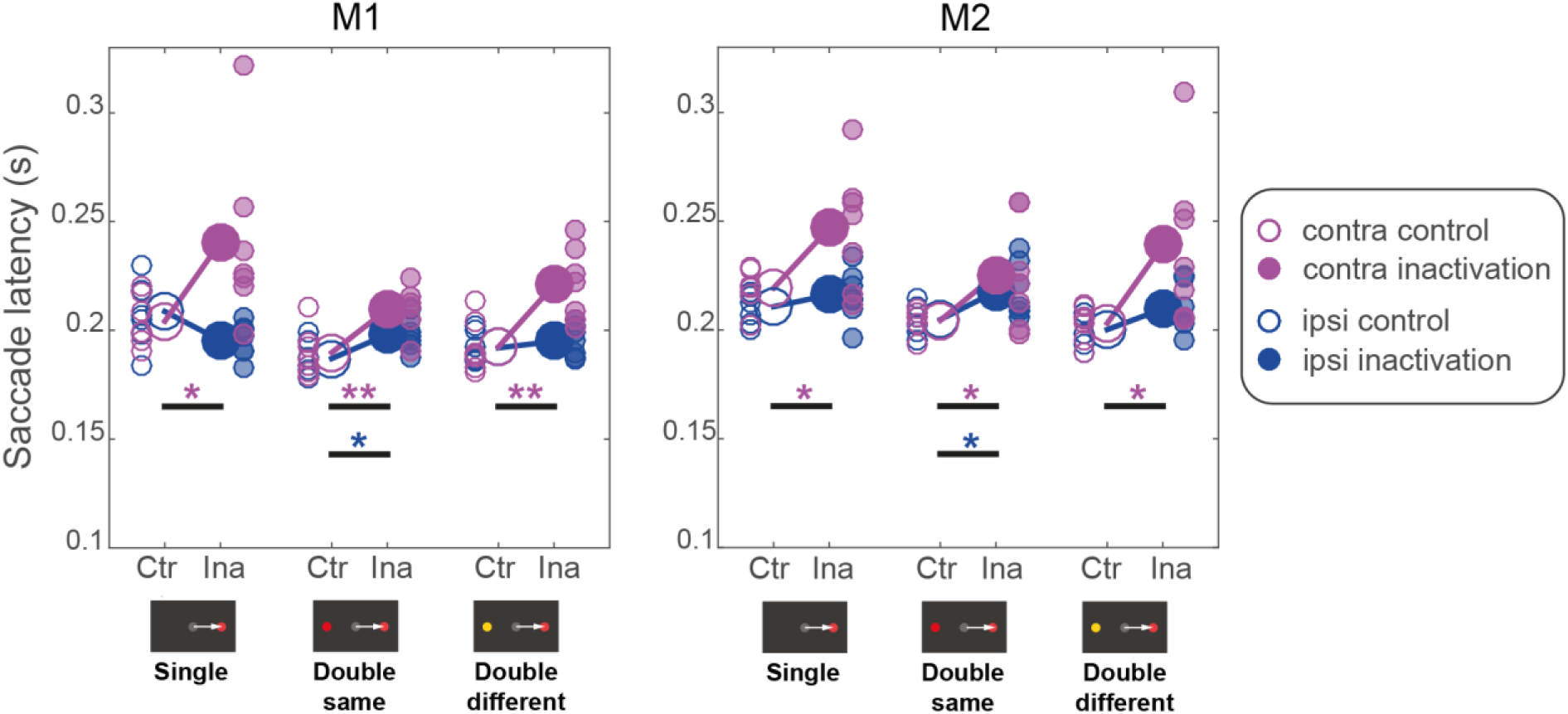
Inactivation effects on saccade latency to the target. The saccade latency is displayed separately for control (empty circles, “Ctr”) and inactivation (filled circles, “Ina”) sessions for each stimulus type and hemifield (icons below horizontal axis show example stimulus display for one hemifield). Small circles display single sessions; large circles display the mean across sessions. We tested the difference between control and inactivation saccade latency of selecting either contralesional (magenta) or ipsilesional (blue) stimuli (independent t-test, one star *, p < .05; two stars **, p<0.01).

**Table 1.** Results of the independent t-test for the comparison of saccade latency in control vs inactivation sessions. The significant effects are in bold font.

| Stimulus type | Hemifield | Monkey 1 (M1) |  | Monkey 2 (M2) |  |
| --- | --- | --- | --- | --- | --- |
|  |  | t-value | p-value | t-value | p-value |
| Single stimuli | **contra** | **2.31** | **.039** | **2.47** | **.029** |
|  | ipsi | 2.07 | .061 | 0.99 | .342 |
| Double same stimuli | **contra** | **3.21** | **.008** | **2.19** | **.049** |
|  | **ipsi** | **2.81** | **.016** | **2.43** | **.032** |
| Double different stimuli | **contra** | **3.62** | **.004** | **2.54** | **.026** |
|  | ipsi | 0.80 | .44 | 1.91 | .081 |

### The effects of stimulus type and inactivation on accuracy

The apparent difference between accuracy for easy vs difficult discrimination in the control sessions (**Figure 3**) was the intended consequence of our task design (since the accuracy level was manipulated experimentally by adjusting the distractor difficulty, we do not present the corresponding statistical comparison). However, when we analyzed each difficulty level separately, we also observed accuracy differences between the three stimulus types during difficult discrimination (repeated measures one-way ANOVA, within-factor “Stimulus Type”; for the difficult discrimination: monkey M1: F(2,20) = 132.26, p < .001; monkey M2: F(2,20) = 115.96, p < .001; for the easy discrimination: M1: F(2,20) = 2.23, p = .15; M2: F(2,20) = 2.54, p = .12). In both monkeys, the accuracy was highest for the double different stimuli and lowest in the double same stimuli for the difficult distractor (post-hoc tests; **Suppl. Table S2**). This difference in accuracy suggests that the three stimulus types elicited different behavioral strategies (for instance, direct comparison between hemifields for double different stimuli vs. only memorized representations of targets and distractors in double same and single stimuli; see later).

**Figure 3.**
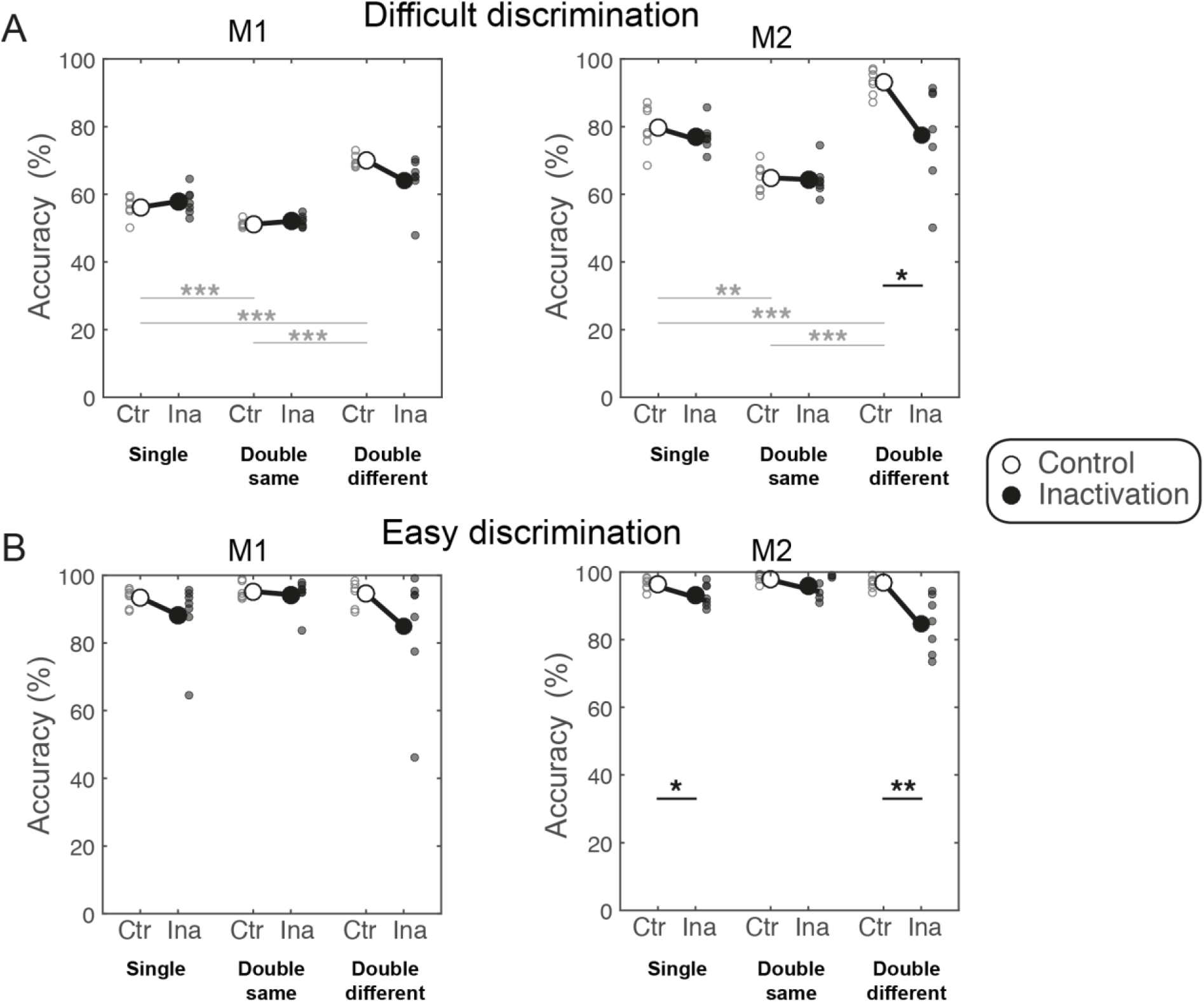
Inactivation effects on accuracy. The accuracy is displayed separately for control (“Ctr”) and inactivation (“Ina”) sessions for each stimulus type and difficulty level. Small circles depict single sessions, and large circles indicate the mean across sessions. Two statistical analyses are presented: the difference between control and inactivation sessions (black connecting lines and stars) and the difference in accuracy between stimulus types (single, double same, double different) for the control sessions (gray connecting lines and stars). (A) Difficult discrimination. The inactivation did not affect the accuracy, besides a decrease in the double different condition in M2. Considering only the control sessions, the accuracy significantly varied between stimulus types. (B) Easy discrimination. The inactivation affected accuracy for M2 for single and double different conditions. The accuracy in the control condition was very high and did not vary between stimulus types. Independent t-test, one star *, p < .05; two stars **, p<0.01; three stars ***, p<0.001.

We next analyzed the effects of inactivation on accuracy using a three-way mixed ANOVA with within-factors “Stimulus Type” (Single / Double Same / Double Different) and “Difficulty” (Difficult / Easy discrimination) and between-factor “Perturbation” (Control / Inactivation sessions). This analysis (see details in **Suppl. Table S3**) revealed no main effect of the factor “Perturbation” and no statistically significant interactions with this factor for M1. However, the main effect of the “Perturbation” and the two-way interaction between Stimulus Type × Perturbation was significant for M2 (F(1,12) = 9.4, p = .01, F(2,24) = 6.4, p = .006).

The results for M2 were followed up to investigate the effect of inactivation by applying post-hoc tests. M2 showed a significant decrease in accuracy for single stimuli for the easy distractor (t(1,12) = −2.26, p = .04) and for double different stimuli for both difficulty levels (difficult: t(1,12) = −2.67, p = .02; easy: t(1,12) = −3.73, p = .003) but not for double same stimuli (difficult: t(1,12) =0.18, p = .86; easy: t(1,12) =1.48, p = .16) or difficult single stimuli (t(1,12) = 0.86, p = .4) (**Figure 3**). These effects will be addressed with the Signal Detection Theory analysis below.

### The effect of inactivation on criterion and d-prime

We adopted the Signal Detection Theory approach to differentiate between the spatial selection bias by calculating the response criterion and the deficit in perceptual discrimination between stimuli by calculating d-prime. To assess at first all possible interaction effects on criterion and d-prime, a four-factor mixed ANOVA was used per monkey, including within-factors “Stimulus Type” (Single / Double Same / Double different), “Difficulty” (Difficult / Easy discrimination) and “Hemifield” (Contra- / Ipsilesional) and between-factor “Perturbation” (Control / Inactivation sessions). This ANOVA revealed that there was a significant interaction of all four factors (Stimulus Type × Difficulty × Hemifield × Perturbation) only in M1 for the d-prime (F(2,24) = 3.87, p = .035) but not for the criterion and not in M2 (criterion: F(2,24) = 0.33, p = .72, d-prime: F(2,24) = 0.33, p = .72). Related to the perturbation, for the criterion M1 showed a three-way Stimulus Type × Difficulty × Perturbation interaction (F(2,24) = 8.69, p = .001), and two two-way interactions (Perturbation × Difficulty, F(1,12) = 17.86, p = .001 and Perturbation × Stimulus Type, F(2,24) = 17.30, p < .001); and for the d-prime, a three-way Perturbation × Hemifield × Stimulus Type interaction (F(2,24) = 3.99, p = .032). Likewise, M2 showed two two-way interactions (Perturbation × Stimulus Type, F(2,24) = 7.82, p = .002; Perturbation × Hemifield, F(2,24) = 6.71, p = .024) for the criterion; and for the d-prime, a main effect of Perturbation (F(1,12) = 10.00, p = .008) and a two-way Stimulus Type × Perturbation interaction (F(2,24) = 8.76, p = .001) (see the details in the in **Suppl. Table S4**).

Although the four-factor mixed ANOVA includes all possible interactions, it is difficult to interpret and it cannot directly answer our research question, such as whether dPul inactivation affects the criterion or d-prime, differently for the three stimulus types and the two perceptual difficulty levels. In both monkeys, we observed interactions of the factors “Perturbation” and “Hemifield”, and we had *a priori* hemifield-specific predictions. To test these predictions, in the sections below we continue the analysis of d-prime and criterion focusing on a two-factor mixed ANOVA (see the details in the **Suppl. Table S5** and **S6**) with the within-factor “Hemifield” (Contra- / Ipsilesional) and the between-factor “Perturbation” (Control / Inactivation sessions), plus the corresponding post-hoc t-tests, separately for each stimulus type and difficulty.

### The effect of inactivation for single stimuli

The single stimuli condition involves a perceptual judgment between making a saccade to a peripheral target or continuing fixating as the correct response to a peripheral distractor (low spatial competition between a central “stay” and peripheral “go” options). To evaluate a deficit in discrimination versus a spatial selection bias, we divided trials into hits, misses, correct rejections, and false alarms separately for stimuli presented in the contralesional hemifield (opposite to the side of inactivation) or ipsilesional hemifield to calculate the hit rate, false alarm rate, d-prime and criterion for each hemifield (**Suppl. Figure S2**). Since we used two distractors: one easy distractor (yellow) that was perceptually clearly different from the red target and the other (orange) distractor required a difficult perceptual discrimination, we analyzed these two distractor conditions separately, contrasting them to target trials. Here and in the next sections, we first describe quantitative predictions using simulated data and then the actual data separately for difficult and easy discrimination.

During difficult discrimination, if dorsal pulvinar inactivation causes a spatial selection bias, we expect a similar decrease in contralesional hit rate and false alarm rate, resulting in a criterion shift towards “less contra” (**Figure 4A**). If the inactivation causes a contralesional perceptual discrimination deficit, we expect a decrease in the contralesional hit rate and an increase in the false alarm rate, resulting in decreased contralesional d-prime (**Figure 4A**). We did not expect changes to the ipsilesional criterion and d-prime.

**Figure 4.**
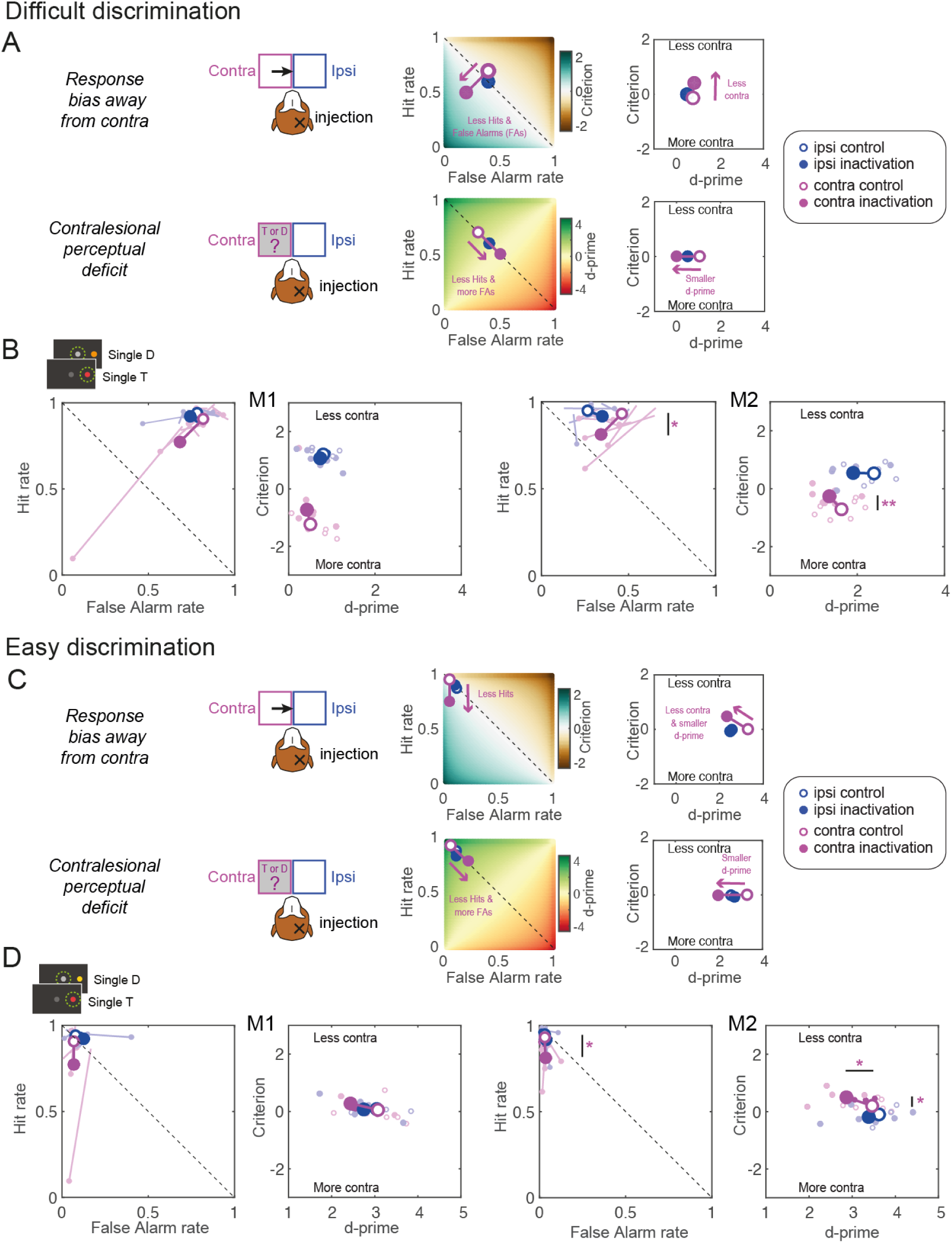
Predictions and results for single stimuli. (A) Illustration of the two alternative hypotheses for the difficult discrimination with simulated data showing the expected changes in hit rate, false alarm rate, d-prime and criterion after unilateral dPul inactivation. The color-coded background in the hit rate vs false alarm rate plots shows the corresponding criterion or d-prime. Positive shifts of the criterion are defined as towards “Less contra” (and vice versa). (B) Inactivation effects on signal detection variables for the difficult discrimination (orange distractor); the data are displayed separately for each monkey, the left panel shows ipsilesional (blue) and contralesional (magenta) false alarm rate and hit rate, the right panel shows the ipsilesional and contralesional criterion and d-prime. Small circles denote single sessions; large circles –mean across sessions. (C) Illustration of the two alternative hypotheses for the easy discrimination. (D) Inactivation effects on signal detection variables for the easy discrimination (yellow distractor). Abbreviations: T – target, D – distractor, contra – contralesional, ipsi – ipsilesional.

In the target trials, the contralesional hit rate decreased after the inactivation significantly for M2 (independent t-test; M1: t(1,12) = 1.1, p = .29; M2: t(1,12) = −2.89, p = .01). For difficult discrimination (**Figure 4B**), the contralesional false alarm rate also decreased, but this effect did not reach significance (M1: t(1,12) = 1.15, p = .27; M2: t(1,12) = 1.6, p = .14). The two-way mixed-effect ANOVA with factors “Perturbation” and “Hemifield” performed on d-prime and criterion showed no significant interaction of “Perturbation” × “Hemifield” in either monkey, and only a significant main effect of “Perturbation” in M2 on criterion (F(1,12) = 6.40, p = .026; see **Suppl. Table S5**). Accordingly, M2 showed a significant shift of the contralesional criterion towards “less contra” (M2: t(1,12) = 3.07, p = .01; the effect was similar but did not reach significance in M1 (t(1,12) = 1.21, p = .25). Neither monkey showed a decrease in contralesional d-prime after inactivation (M1: t(1,12) = −0.36, p = .73; M2: t(1,12) = −1.26, p = .23). In line with our predictions, in both monkeys neither ipsilesional d-prime nor the ipsilesional criterion exhibited any changes (d-prime M1: t(1,12) = −0.41, p = .69; M2: t(1,12) = −1.98, p = .07; criterion M1: t(1,12) = −0.99, p = .34; M2: t(1,12) = 0.15, p = .89).

During easy discrimination, if dorsal pulvinar inactivation causes a spatial selection bias, we expect a decrease in the contralesional hit rate but now no change in false alarm rate due to a “floor effect” (already very low false alarm rate in the control sessions). This will result in a shift of criterion towards “less contra” combined with a decrease in contralesional d-prime (**Figure 4C**). For completeness, if the inactivation causes a contralesional perceptual discrimination deficit, one expects a decrease in contralesional hit rate and an increase in false alarm rate (although given the easy discriminability of the yellow distractor, we did not expect such an increase in the actual behavior). This would result in decreased contralesional d-prime but no change in criterion (**Figure 4C**).

Indeed, in both monkeys during easy discrimination (**Figure 4D**), the false alarm rate was already near zero, so there was no room to exhibit any inactivation-induced decrease (independent t-test; contra M1: t(1,12) = 0.02, p = .98; M2: t(1,12) = −0.3, p = .7; ipsi M1: t(1,12) = 0.95, p = .36; M2: t(1,12) = −0.6, p = .6). In a two-way mixed-effect ANOVA on d-prime and criterion, there was no significant main effect for “Perturbation” or any interaction between “Perturbation” and “Hemifield” in both monkeys (in M2, the interaction for the criterion showed a trend, F(1,12) = 4.59, p = .053, see **Suppl. Table S6**). Accordingly, M2 showed a significant shift for the contralesional criterion towards “less contra” (M1: t(1,12) = 0.85, p = .41; M2: t(1,12) = 2.63, p = .02), but also a decrease in contralesional d-prime (M1: t(1,12) = −1.49, p = .16; M2: t(1,12) = 2.27, p = .04). This decrease in d-prime was due to the “floor effect” of already very low false alarm rate, as predicted in the simulation.

To sum up, under conditions of low spatial competition, after inactivation both monkeys showed a shift in response criterion manifesting as reluctance to select stimuli in the contralesional hemifield for both difficulty levels, significant in monkey M2.

### The effect of inactivation for double same stimuli

Previous studies suggested that dorsal pulvinar becomes most relevant in the case of spatial competition between hemifields (Desimone et al., 1990; Wilke et al., 2013; Dominguez-Vargas et al., 2017). Here, two equally rewarded targets or two distractors were presented in the periphery during the double same stimuli condition, eliciting a high competition between hemifields for visual representation and response selection. The monkeys chose between continuing fixating or making a saccade to one of the two peripheral stimuli (**Suppl. Figure S3**).

As for the single stimuli, if dorsal pulvinar inactivation causes a spatial selection bias during inter-hemifield competition, we expect a similar decrease in contralesional hit and false alarm rates. Necessarily, such a decrease has to result in either (i) a corresponding increase in ipsilesional hit rate and false alarm rate, or (ii) an increase of central fixation selection. Both monkeys tended to select peripheral stimuli over the central fixation, even after inactivation, as shown with a non-hemifield-selective criterion analysis (“stay” vs. “go”, **Suppl. Table S7**). Therefore, we expected a shift for both contralesional and ipsilesional criteria towards “less contra” (**Figure 5A**). If dorsal pulvinar inactivation causes a contralesional perceptual discrimination deficit, similarly to the single stimuli condition we expect a decrease in contralesional hit rate and an increase in contralesional false alarm rate, resulting in a reduction of contralesional d-prime (**Figure 5A**). We did not have strong predictions for the ipsilesional discrimination: it might remain unaffected, as for single stimuli (and instead, only fixation selection might change to counterbalance the contralesional changes, the possibility we illustrate here), or it might improve as a consequence of ipsilesional hit rate increase and ipsilesional false alarm rate decrease.

**Figure 5.**
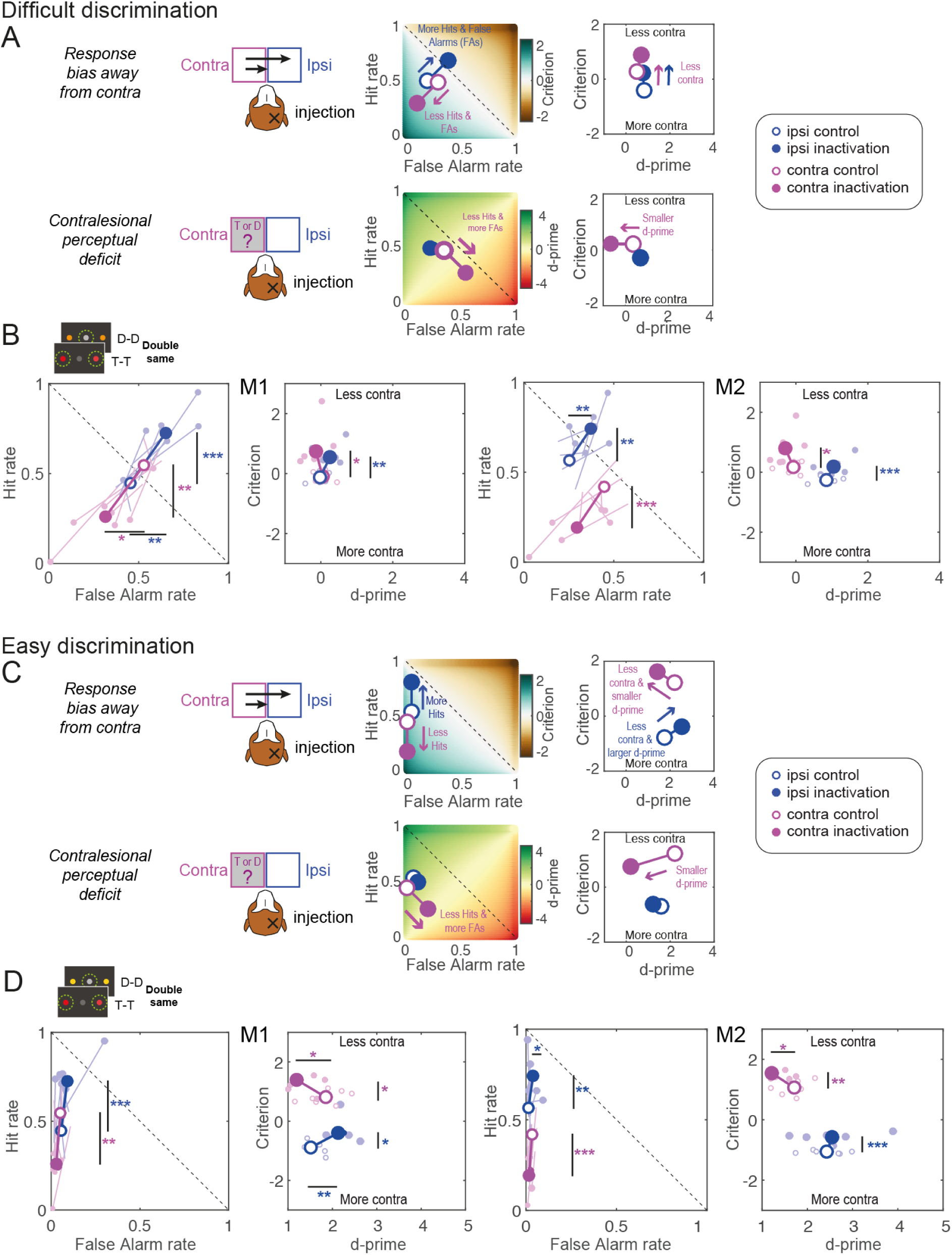
Predictions and results for double same stimuli. Same format and notations as in Figure 4. (A) Illustration of the two alternative hypotheses for the difficult discrimination in case of the double same stimuli. (B) Inactivation effects on signal detection variables for the difficult discrimination, with data separately shown for each monkey, with single sessions and overall means displayed. (C) Illustration of the two alternative hypotheses for the easy discrimination in case of the double same stimuli. (D) Inactivation effects on signal detection variables for the easy discrimination. Abbreviation: T – target, D – distractor, contra – contralesional, ipsi – ipsilesional.

The contralesional hit rate decreased significantly for both monkeys (independent t-test; M1: t(1,12) = −4.24, p < .001; M2: t(1,12) = −4.78, p < .001), and ipsilesional hit rate increased significantly (M1: t(1,12) = 4.36, p < .001; M2: t(1,12) = 3.28, p = .01). For difficult discrimination displayed in **Figure 5B**, the ipsilesional false alarm rate significantly increased for both monkeys (M1: t(1,12) = 3.25, p = .01; M2: t(1,12) = 3.23, p = .01), and contralesional false alarm rate significantly decreased for M1 (M1: t(1,12) = −2.79, p = .02; M2: t(1,12) = −2.04, p = .065). The two-way mixed-effect ANOVA performed on d-prime and criterion showed no significant “Perturbation” × “Hemifield” interaction (see **Suppl. Table S5**). For the criterion, a significant main effect for “Perturbation” was observed for both monkeys (M1: F(1,12) = 9.65, p = .009; M2: F(1,12) = 10.43, p = .007). Both monkeys showed a significant shift of the contralesional and ipsilesional criterion towards “less contra” (M1 contra: t(1,12) = 2.68, p = .02, ipsi: t(1,12) = 3.74, p = .003; M2 contra: t(1,12) = 2.87, p =.014, ipsi: t(1,12) = 3.8, p = .003). Neither monkey showed a significant change in contralesional d-prime (p > .05).

During easy discrimination, as for the single stimuli, for the spatial selection bias hypothesis we expect a decrease in contralesional hit rate and no observable decrease in false alarm rate (again due to the “floor effect” on already very low false alarm rate), and a corresponding increase in ipsilesional hit rate, but no increase in false alarm rate, due to easy discriminability of the distractor. These changes should result in a shift for both contralesional and ipsilesional criteria towards “less contra” combined with a change in d-prime values (**Figure 5C**). For the contralesional perceptual discrimination deficit hypothesis, we expect a decrease in the contralesional hit rate and an increase in the false alarm rate resulting in a decrease in contralesional d-prime (**Figure 5C**).

For the criterion, we found a significant main effect for “Perturbation” in both monkeys (M1: t(1,12) = 8.30, p = .014; M2: t(1,12) = 31.20, p < .001). For the d-prime, we found significant interaction of “Perturbation” × “Hemifield” in M1 (F(1,12) = 11.99, p = .005; **Suppl. Table S6**). To further evaluate the selection behavior for easy discrimination, we examined the results of the follow-up tests and compared them with the hypothesis-driven simulations. For easy discrimination (**Figure 5D**), the false alarm rate was already near zero, so there was no room to exhibit any inactivation-induced decrease: the contralesional false alarm rate did not show an effect (independent t-test; M1: t(1,12) = −0.94, p = .2; M2: t(1,12) = 1.82, p = .09). The ipsilesional false alarm rate slightly increased, significant for one monkey (M1: t(1,12) = 1.37, p =0.37; M2: t(1,12) = 2.32, p= .04). Consequently, both monkeys showed a significant shift for the contralesional and ipsilesional criterion towards “less contra” (M1 contra: t(1,12) = 2.83, p = .015, ipsi: t(1,12) = 2.66, p = .021; M2 contra: t(1,12) = 3.6, p = .004, ipsi: t(1,12) = 7.66, p < .001). Both monkeys also showed a significant decrease in contralesional d-prime (M1: t(1,12) = −2.32, p = .039; M2: t(1,12) = −2.30, p = .04), due to the already very low false alarm rate, and M1 showed an increase in ipsilesional d-prime (M1: (1,12) = 3.31, p = .006, M2: (1,12) = −0.38, p = .71), due to increase in ipsilesional hit rate but without corresponding increase in ipsilesional false alarm rate. Overall, the data for both difficulty levels are consistent with the spatial selection bias hypothesis.

### The effect of inactivation for double different stimuli

Similar to the double same stimuli, the double different stimuli condition also comprises a high spatial competition between hemifields. Furthermore, might be influenced by the possibility of directly comparing the simultaneously presented target and distractor in the opposite hemifields. Notably, the central fixation is always an incorrect response option in this condition (**Suppl. Figure S4**).

During difficult discrimination, for the spatial selection bias hypothesis we expect the same effects as for the double same stimuli (**Figure 6A**). But for the perceptual discrimination deficit hypothesis and assuming the “go” bias, in contrast to double same stimuli here we expect the decrease in contralesional d-prime to be necessarily linked to the *decrease* in ipsilesional d-prime. This is because selecting less targets on the contralesional side would lead to selecting *more* distractors on the ipsilesional side, and selecting more contralesional distractors – to *less* ipsilesional targets (**Figure 6A**).

**Figure 6.**
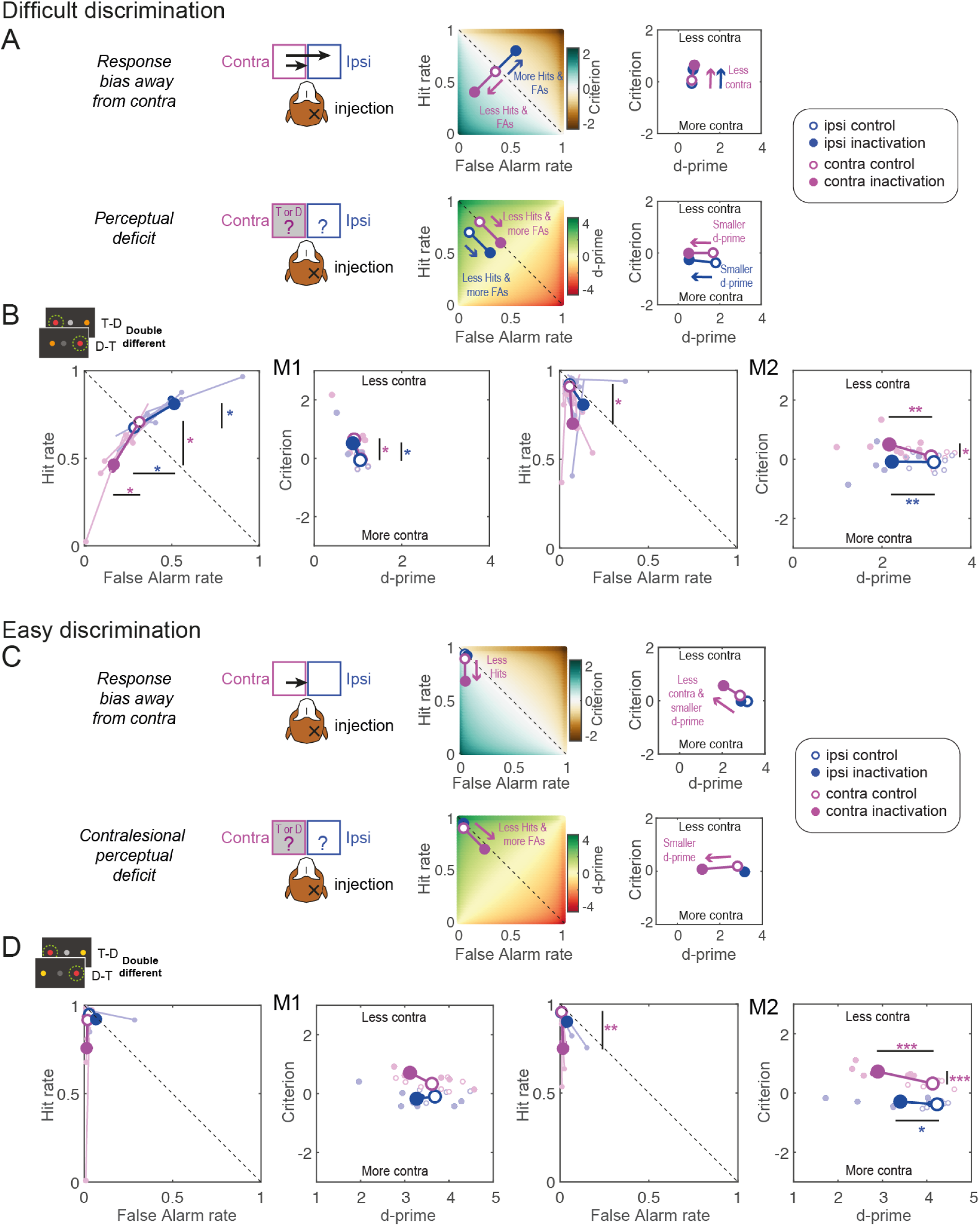
Predictions and results for double different stimuli. Same format and notations as in Figure 4. (A) Illustration of the two alternative hypotheses for the difficult discrimination in case of the double different stimuli. (B) Inactivation effects on signal detection variables for the difficult discrimination, with data separately shown for each monkey, with single sessions and overall means displayed. (C) Illustration of the two alternative hypotheses for the easy discrimination in case of the double different stimuli. (D) Inactivation effects on signal detection variables for the easy discrimination. Abbreviation: T – target, D – distractor, contra – contralesional, ipsi – ipsilesional.

During difficult discrimination, the contralesional hit rate decreased significantly in both monkeys (independent t-test; M1: t(1,12) = −2.8, p = .02; M2: t(1,12) = −2.98, p = .01) and ipsilesional hit rate increased for M1 (M1: t(1,12)= 2.92, p = .01; M2: t(1,12) = 1.49, p = .16; **Figure 6B**). The contralesional and ipsilesional false alarm rate significantly decreased only for M1 (contra: M1: t(1,12) = −2.9, p = .01; M2: t(1,12) = −0.65, p = .53; ipsi: M1: t(1,12) = −2.9, p = .01; M2: t(1,12) = −1.8, p = .09). Consequently, we observed for the criterion a significant main effect of “Perturbation” in M1 (F(1,12) = 6.47, p = .026) and an interaction of “Perturbation” × “Hemifield” in M2 (M2: F(1,12) = 6.88, p = .022; **Suppl. Table S5**). In line with the response bias hypothesis, M1 showed a significant shift for the contralesional and ipsilesional criterion towards “less contra” (contra: t(1,12) = 2.4, p = .03, ipsi: t(1,12) = 2.72, p = .02) and no effect for contralesional or ipsilesional d-prime (contra: t(1,12) = −1.42, p = .18, ipsi: t(1,12) = −2.03, p = .06). Likewise, M2 showed a significant shift for the contralesional criterion towards “less contra” (M2: t(1,12) = 2.42, p = .03). But, in accordance with a significant main effect of “Perturbation” for the d-prime in M2 (F(1,12) = 10.73, p = .007), M2 also exhibited a significant decrease in contralesional and ipsilesional d-prime (contra: t(1,12) = −3.14, p = .01; ipsi: t(1,12) = −3.32, p = .006).

During easy discrimination in the presence of a yellow distractor, for the spatial selection bias hypothesis, we expect a decrease in a contralesional hit without the decrease in false alarm rate due to the “floor effect”. We also expect no increase in the ipsilesional hit rate because it is already very high and no increase in the ipsilesional false alarm rate because of the easy discriminability of the yellow distractor (**Figure 6C**). For contralesional perceptual discrimination deficit, similarly to the double same stimuli, we expect a decrease in contralesional d-prime but no effect on ipsilesional d-prime (**Figure 6C**).

For easy discrimination (**Figure 6D**), the false alarm rate was already near zero, so there was no room to exhibit any inactivation-induced contralesional decrease (independent t-test; M1: t(1,12) = 0.8, p = .47; M2: t(1,12) = −1.87, p = .09). The ipsilesional false alarm rate did not increase (M1: t(1,12) = 0.98, p = .35; M2: t(1,12) = 1.66, p = .12). The contralesional hit rate decreased for M2 (M1: t(1,12) = −1.2, p = .25; M2: t(1,12) = −4.22, p < .001). In the ANOVA, for the criterion we found a significant interaction of “Perturbation” × “Hemifield”, and the main effect of “Perturbation” in M2 but not M1 (F(1,12) = 9.52, p = .009; F(1,12) = 22.96, p < .001; **Suppl. Table S6**). Accordingly, only M2 showed a significant shift for the contralesional criterion towards “less contra” (M1: t(1,12) = 0.82, p = .25; M2: t(1,12) = 4.75, p < .001). Furthermore, in M2 there was a significant main effect of “Perturbation” for the d-prime (F(1,12) = 13.49, p = .003) and a corresponding decrease in contralesional and ipsilesional d-prime (contra M1: t(1,12) = −0.82, p = .43; M2: t(1,12) = −5.16, p < .001; ipsi: M1: t(1,12) = −2.03, p = .07; M2: t(1,12) = −2.19, p = .049). This decrease in d-prime was due to the “floor effect” of already very low false alarm rate.

To sum up these results, the inactivation effects for M1 during difficult discrimination fully matched the predictions of the spatial selection bias hypothesis (shift of both contralesional and ipsilesional criteria towards “less contra”). In contrast, in M2, only the contralesional criterion, for both difficulty levels, shifted towards “less contra”. All stimulus conditions together are summarized in **Table 2** and **Suppl. Figures S5** and **S6**. For M2, we also observed a decrease in contralesional d-prime for both difficulty levels and a decrease in ipsilesional d-prime for the difficult discrimination. The contralesional d-prime decline can be accounted for by the already very low false alarm rate (for both difficulty levels), as in the single stimuli and double same stimuli conditions for easy discrimination. But the ipsilesional d-prime decrease, which only manifested in the double different stimuli condition, can be related to the compromised ability of this monkey to utilize the information from the contralesional hemifield for the direct comparison with the ipsilesional stimulus – the strategy that the monkey successfully relied on in the control sessions (*cf.* **Figure 3**). We further address this proposition in the Discussion.

**Table 2.**
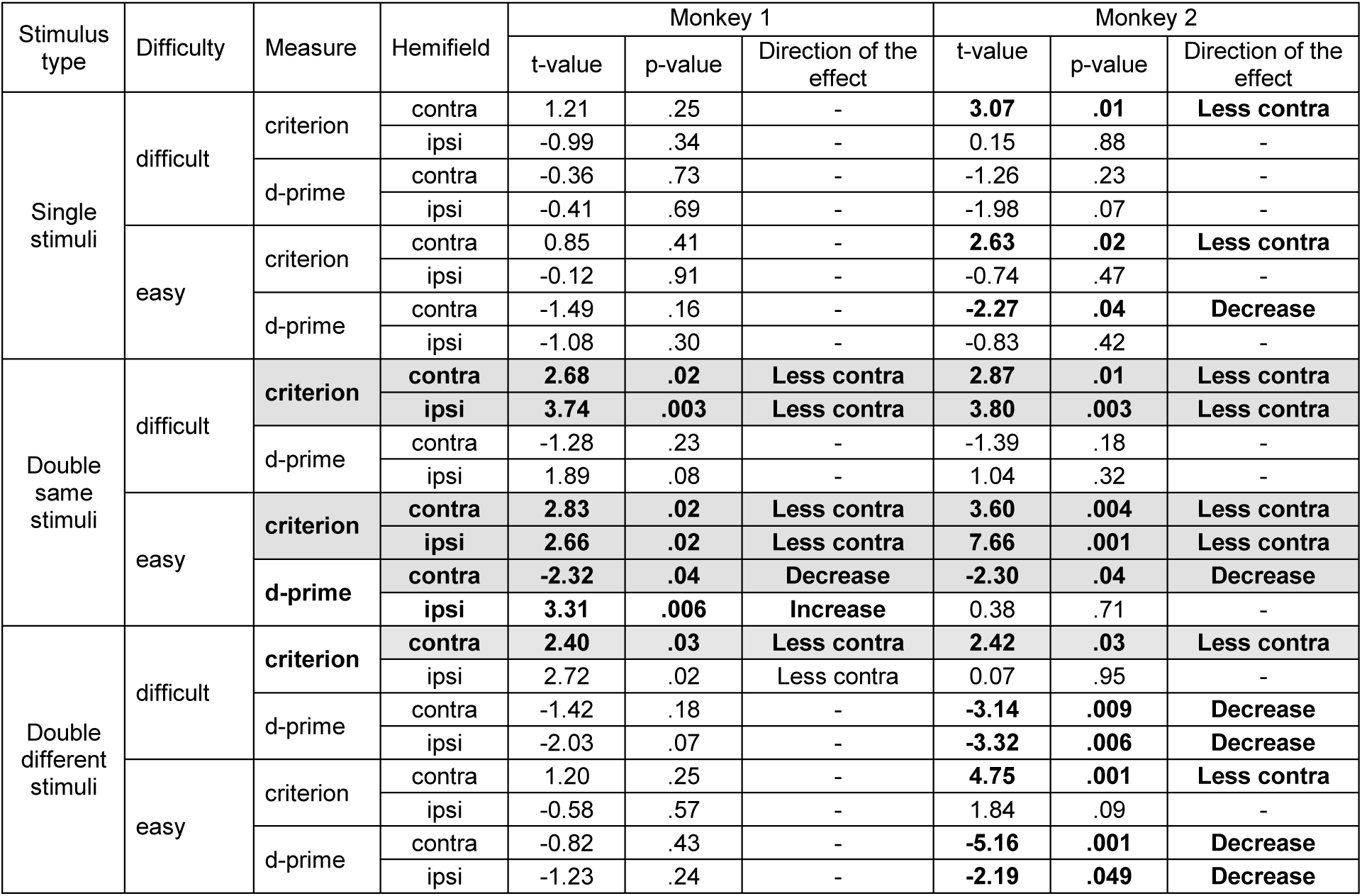
Summary of inactivation effects for the criterion and d-prime using independent t-tests. Significant effects are in bold font, consistent effects across the two monkeys are highlighted with gray background. **Suppl. Figures S5** and **S6** plot the summary of the corresponding data, and **Suppl. Table S8** for nonparametric tests.

## Discussion

This study utilized task demands such as fast perceptual color discrimination between target and distractor, spatially competing stimuli, and stimulus-congruent saccade responses to investigate whether the impairments in contralesional perceptual discrimination, as opposed to more general response bias, might contribute to visuospatial deficits after dorsal pulvinar inactivation. Following the inactivation, we primarily observed slowing of contralesional saccades and criterion shifts away from contralesional stimuli, especially when two peripheral stimuli elicited high spatial competition between hemifields (**Table 2**). These effects were present at both perceptual difficulty levels. Notably, the d-prime and the overall accuracy remained largely unaffected. We conclude that the contralesional visuospatial deficits observed after inactivating the dorsal pulvinar are not caused by a contralesional perceptual deficiency but by a spatial selection bias, even during a demanding perceptual decision task.

We adopted the Signal Detection Theory approach to differentiate between spatial selection bias and discrimination sensitivity by calculating the response criterion and d-prime, as has been done in studies of visual and prefrontal cortices (Luo and Maunsell, 2015, 2018) and the superior colliculus (McPeek and Keller, 2004; Lovejoy and Krauzlis, 2017; Crapse et al., 2018). After pulvinar inactivation, a shift in criterion manifested as reluctance to select stimuli in the contralesional hemifield, regardless of whether it contained a target or a distractor. This spatial selection bias, and delays in making a saccade to contralesional stimuli, were observed for all three stimulus conditions (single, double same, and double different). In our experiments, as in many prior studies, the saccadic “report” coincided with the target stimulus location. Therefore, the observed change in the decision criterion could be attributed to both a (pre)motor “response bias”, as well to a more stimulus position-specific “choice bias” that would also be revealed by a dissociated saccade or even a manual report (Sridharan et al., 2017; Crapse et al., 2018; Luo and Maunsell, 2019). Note however that we purposely designed the task to probe pulvinar under ecologically-valid conditions where sensory information aligns with the spatial goals.

Similar to our previous work (Wilke et al., 2010, 2013), here we found a pronounced selection bias “away from contra” for the double same stimulus condition most similar to the free-choice between two identical targets. But also for single stimuli, the shift of the criterion “away from contra” was significant for one monkey and similar (but not significant) for the other monkey. In contrast to the current results, Wilke et al. (2010, 2013) did not find a decrease in the correct selection of contralesional single instructed targets. In these prior studies, the instructed single target condition differed from the current single stimulus condition because monkeys invariably had to saccade to the target. Here we included a second response option, staying at the central fixation point, as the correct response for the distractor trials. The presence of two options and the perceptual discrimination task context created a low level of spatial competition between the fixation and the peripheral option in the single stimuli condition - sufficient to engender an effect from pulvinar inactivation. Hence, the dorsal pulvinar inactivation influenced the competition not only between the hemifields but also the competition between the foveal and the peripheral contralesional options.

Support for the importance of spatial competition also comes from a brief report by Desimone and colleagues, who inactivated the lateral pulvinar in one macaque during a color rule task that required a specific response with a manual lever to a briefly flashed colored target and not to a distractor (Desimone et al., 1990). The target was defined as the stimulus appearing at the same location as the briefly flashed cue. The error rate increased when the cue and the target were in contralesional hemifield while the conflicting distractor was in the ipsilesional hemifield. However, when both the target and the distractor were located within the same hemifield – thereby obviating the competition between hemifields – pulvinar inactivation did not yield any significant impact on performance.

Notably, in the present study the inactivation-induced selection bias occurred even when only one response option was correct and rewarded. Placing a target in the contralesional hemifield when it was the only rewarded option (i.e. single target or target-distractor conditions) did not alleviate the spatial selection bias “away from contra”. Both monkeys selected the contralesional target less and instead chose the ipsilesional distractor or the fixation option, receiving no reward in these trials. Again, this contrasts with the observed alleviation of the selection bias in the value-based saccade choice task where a color cue explicitly and unambiguously signaled a large reward (Wilke et al., 2013). Another difference to the present study was that in Wilke et al. there was a memory delay that monkeys could have utilized to make a deliberate value-based choice. The present results emphasize the contribution of dPul to fast decisions between conflicting options under uncertainty, in agreement with the previous inactivation study in the lateral pulvinar (Desimone et al., 1990), and in agreement with a microstimulation study from our group which showed a strong microstimulation-induced choice bias in the immediate visually-guided saccade task but not in the delayed memory-guided task (Dominguez-Vargas et al., 2017).

A hypothetical inactivation-induced contralesional perceptual discrimination deficit should engender to a confusion between targets and distractors in the contralesional hemifield. This would lead to an increased false alarm rate alongside a decreased hit rate, resulting in a decreased contralesional d-prime. We did not observe a pattern that would be consistent with a contralesional perceptual discrimination deficit in any of the three stimulus type conditions. Likewise, previous pulvinar lesion studies in non-human primates showed unimpaired visual discrimination learning (Ungerleider and Christensen, 1977; Bender and Butter, 1987) and unimpaired contralesional visual motion discrimination performance without spatially-competing distractors (Komura et al., 2013). These studies however used manual responses that were spatially dissociated from the visual stimuli. Our study extends these findings to situations in which saccadic responses are spatially-contingent on the stimuli, and there is a competition between spatial locations. But even under these conditions, where the contribution of the pulvinar might be more crucial, the perceptual sensitivity was largely unaffected after pulvinar inactivation.

One possible strategy to accurately discriminate targets from distractors in our task is to compare the presently visible stimuli with learned and memorized representations of the target and the distractors. This strategy could be used for all three stimulus types. For the double different stimuli condition, however, an additional strategy could be employed. The visual appearance of the two stimuli, one target, and another distractor, could be directly compared across hemifields without relying upon, or in addition to, memorized representations. Indeed, both monkeys had significantly higher accuracy in the target-distractor condition than in the single or double same stimuli conditions for the difficult discrimination. Furthermore, the accuracy in easy and difficult target-distractor trials was very high and did not significantly differ during control sessions in monkey M2. Hence, the strategy based on the direct comparison improved discrimination performance for the target-distractor condition.

After inactivation, M2’s accuracy decreased substantially for the difficult target-distractor condition, driven by a significant drop in both, contralesional and ipsilesional d-prime. We argue that this is not an indication of a specific contralesional perceptual discrimination deficit but a consequence of a direct across-hemifield comparison strategy, and its partial failure. In this interpretation, M2 avoided going to the contralesional hemifield (due to the criterion shift), but still could utilize the information from the ipsilateral hemifield to correctly reject contralesional distractors, as demonstrated by a very low contralesional false alarm rate even after the inactivation. The resulting contralesional sensitivity decrease is thus similar to the case of easy discrimination but the ipsilesional hit rate did not increase (on the contrary, it decreased non-significantly). Instead, the contralesional hit rate decrease was compensated by more frequent central fixations. Furthermore, the ipsilesional false alarm rate increased non-significantly. We suggest that the ipsilesional sensitivity decreased because the inactivation disrupted the access to information from contralesional hemifield for comparing it to the stimulus in the ipsilesional hemifield.

Pulvinar lesions in humans (Karnath et al., 2002; Arend et al., 2008a; Snow et al., 2009) and monkeys (Petersen et al., 1987; Desimone et al., 1990; Robinson and Petersen, 1992; Zhou et al., 2016) lead to deficits in spatial attention tasks. Prior work also showed that subjects may shift either their criterion or sensitivity at the attended location relative to the unattended location (Wyart et al., 2012; Luo and Maunsell, 2015, 2018). These studies raise the possibility of relating our findings to visual spatial attention. Experiments investigating spatial attention typically use a valid or invalid cue indicating where to attend to an upcoming target without making an eye movement (Posner et al., 1980; Petersen et al., 1987; Lovejoy and Krauzlis, 2010; Luo and Maunsell, 2015, 2019; Fiebelkorn et al., 2019). Our task was not designed to investigate covert spatial attentional processes or shifts of attention, as it lacked the attentional cue. Nevertheless, our main finding of the spatial selection bias after dorsal pulvinar inactivation generally fits well with the previous work emphasizing the role of the pulvinar in selective spatial attention (Kastner and Pinsk, 2004; Halassa and Kastner, 2017).

Indeed, spatial choice bias is often considered a component, or a manifestation of attentional processing, because it captures the competition between spatial locations (Krauzlis et al., 2014; Jaramillo et al., 2019). Desimone and Duncan proposed interpreting the findings after the unilateral pulvinar inactivation as formulated in the biased competition theory (Desimone and Duncan, 1995). In this framework, unilateral inactivation puts the contralesional hemifield at a disadvantage for selection by biasing the ongoing competition for the neuronal representation of multiple stimuli, hence leading to the observed bias “away from contra”. When stimuli compete for attention, attending to one option biases the competition by enhancing the activity in the neurons representing this response option within their receptive field. We speculate that the activity in the pulvinar – and/or in the connected cortical regions – in response to a salient yellow distractor will be suppressed compared to a target or a difficult distractor (target > difficult distractor > easy distractor) – as has been found in V4 during visual search in the presence of a salient pop-out color distractor (Klink et al., 2023). After pulvinar inactivation, we expect a decrease in the saliency (and in the underlying activity) of contralesional stimuli – especially when they are not selected – and corresponding increase in the salience of ipsilesional stimuli due to a push-pull between competing representations (Kinsbourne, 1977). At the same time, we expect that the graded response amplitude for targets and distractors will persist.

Given tight links between attention, perceptual decision and confidence (Kurtz et al., 2017; Denison et al., 2018), the inactivation-induced changes in contralesional salience or criterion might also lead to an altered confidence in choosing the correct response (Fetsch et al., 2014). Our task did not involve confidence judgments. It is plausible however that decreased contralesional confidence in choosing the correct response, operationalized as the increased frequency of opt-out choices (Komura et al., 2013), might result in a spatial selection bias ‘away from contra’ that we have observed. Conversely, the diminished confidence about contralesional stimuli may be a consequence of the observed criterion change. Further studies need to investigate the interplay between response and choice bias, sensitivity, bottom-up or/and top-down saliency of a stimulus, and confidence, by utilizing tasks designed to dissociate between these contributing factors (Wyart et al., 2012; Luo and Maunsell, 2015, 2019; Crapse et al., 2018; Linares et al., 2019).

While in the present study we focused on the dorsal pulvinar, the ventral pulvinar (encompassing the ventral part of the lateral pulvinar, PLvl, and the inferior pulvinar) is also associated with visual attention and salience (Saalmann et al., 2012; Saalmann and Kastner, 2015; Zhou et al., 2016). In particular, the inferior pulvinar can be considered a more perceptually-relevant visual nucleus due to its extensive connectivity to the primary visual cortex and the ventral visual stream (Kaas and Lyon, 2007; Bridge et al., 2016; Kaas and Baldwin, 2020; Kagan et al., 2021). Given such connectivity, visual response properties (Bender, 1981; Petersen et al., 1985; Arcaro et al., 2015, 2018; Zhou et al., 2016), visual target-related microstimulation effects (Dominguez-Vargas et al., 2017), and strong perceptual modulation (Wilke et al., 2009), it would be interesting to investigate if the ventral pulvinar inactivation engenders a more substantial perceptual deficit than the dorsal pulvinar.

In regard to both, the dorsal and the ventral pulvinar, in the human neuroimaging and patient literature the function of pulvinar is sometimes described as distractor filtering (Fischer and Whitney, 2012; Lucas et al., 2019). For instance, LaBerge and Buchsbaum (1990) investigated the contribution of the pulvinar to visual distractor processing using positron emission tomography in healthy subjects. Increased pulvinar activation was found when the target in the contralateral visual field was surrounded by distractors relative to no distractors. The authors concluded that identifying an object in a cluttered visual scene might involve the pulvinar through a filtering mechanism. Similarly, several fMRI studies showed increased activation during more demanding distractor conditions – but these effects could have been driven by task difficulty and response competition rather than specific contralateral filtering processing (Kastner et al., 2004; Strumpf et al., 2013). Likewise, very few patient studies support the notion that the pulvinar participates in filtering out distracting visuospatial inputs in the contralateral hemifield. A study on oculomotor capture found that patients with unilateral pulvinar lesions made slightly more errors when the target was ipsilesional and the distractor was contralesional, compared to the reverse configuration – although the effect was very small (3% error rate difference) (Van der Stigchel et al., 2010). Another study in patients with ventral pulvinar lesions showed impaired contralateral perceptual discrimination in the presence of flanking distractors (Snow et al., 2009).

One interpretation of the distractor filtering hypothesis is that unilateral pulvinar inactivation should disrupt monitoring and filtering out the distractors in the *contralesional* hemifield. If so, we should observe an increased selection of contralesional distractors. We however found no inactivation-induced increase in false alarm rate when an easy or a difficult distractor was presented in the contralesional hemifield. An additional counterargument against the contralateral distractor filtering hypothesis can be derived from cued spatial attention tasks. The inactivation of the lateral pulvinar decreased the error rate due to contralesional distractors – i.e. decreased contralesional false alarm rate (Desimone et al., 1990). Similarly, dorsal lateral pulvinar inactivation caused faster reaction times, compared to control, when the distracting invalid cue was contralesional and the target ipsilesional (Petersen et al., 1987). Collectively, these observations, in conjunction with our findings, challenge the role of the pulvinar in filtering out contralesional distractors, particularly under conditions where stimuli are isolated and not subject to crowding. Instead, the pulvinar might be crucial for the contralateral spatial orienting and selective attention, as was also suggested in a number of patient studies that typically combined dorsal and ventral pulvinar lesions due to stroke etiology (Danziger et al., 2001, 2004; Ward and Danziger, 2005; Ward and Arend, 2007).

Beyond the pulvinar, an ipsilesional selection bias and/or increased contralesional saccade latencies often result from perturbations of frontoparietal cortex and the superior colliculus. For instance, one or both such effects occur after inactivating the lateral intraparietal area LIP during a visual search (Wardak et al., 2002, 2004) and memory saccade tasks (Li et al., 1999; Wilke et al., 2012; Katz et al., 2016; Christopoulos et al., 2018), after inactivating frontal eye fields (FEF) in a visual search task (Wardak et al., 2006), and after parietal or prefrontal cortex inactivation during stimulus-onset asynchrony saccade task (Kubanek et al., 2015; Johnston et al., 2016). These general similarities are in line with dorsal pulvinar’s reciprocal anatomical (Blatt et al., 1990; Hardy and Lynch, 1992; Romanski et al., 1997; Gutierrez et al., 2000) and functional connections (Arcaro et al., 2018; Fiebelkorn et al., 2019; Kagan et al., 2021) with the frontoparietal cortex. In addition to the frontoparietal connectivity, recent work also emphasized the role of the superior temporal regions – which are also strongly interconnected with the pulvinar – in visuospatial processing (Bogadhi et al., 2019). Of course, the specifics of impairments differ between the regions and the paradigms. For instance, the reaction time deficits in visual search target detection depend on the perceptual task difficulty after LIP but not after FEF inactivation (Wardak et al., 2006). In the context of the Signal Detection Theory, while several studies pointed to decision criterion changes after the superior colliculus perturbation (McPeek and Keller, 2004; Sridharan et al., 2017; Crapse et al., 2018), the effect on perceptual sensitivity has also been demonstrated (Lovejoy and Krauzlis, 2017). A major challenge for future research is to compare specific contributions of the pulvinar with other subcortical structures that mediate selective attention and target selection via thalamo-cortical pathways, such as the superior colliculus and basal ganglia (Zenon and Krauzlis, 2012; Crapse et al., 2018; Bogadhi et al., 2019), for example by using pathway-selective manipulations (Klink et al., 2021; Vanduffel and Isa, 2023).

## Methods

### Experimental procedures

All experimental procedures complied with the ARRIVE guidelines (https://arriveguidelines.org) and were conducted in accordance with the European Directive 2010/63/EU, the corresponding German law governing animal welfare, and German Primate Center institutional guidelines. The procedures were approved by the responsible government agency (Niedersaechsisches Landesamt fuer Verbraucherschutz und Lebensmittelsicherheit (LAVES), Oldenburg, Germany).

Two adult male rhesus monkeys (*Macaca mulatta*), weighing 9 kg and 10.5 kg, served as subjects (monkey 1, Cornelius, M1; monkey 2, Curius, M2). For both monkeys, the surgical procedures for implanting an MRI-compatible headpost and chambers, and small within-chamber craniotomies, were the same as described in Dominguez-Vargas et al. (2017). The inactivation locations in the dorsal pulvinar were estimated based on anatomical MRI as described in more detail in the previous work (Dominguez-Vargas et al., 2017; Kagan et al., 2021). Based on the MRI images, we planned where to target the dorsal pulvinar (the grid hole and depth) using the software Planner (Ohayon and Tsao, 2012) and BrainVoyager (Version 2.4.2, 64-bit; Brain Innovation). To confirm the inactivation locations, we performed MRI contrast agent gadolinium injections (**Figure 1A**).

The neural activity was reversibly suppressed using the GABA-A agonist 4,5,6,7-tetrahydro isoxazole [5,4-c]-pyridine-3-ol (THIP hydrochloride; Tocris). The THIP was dissolved in phosphate-buffered saline (PBS). The solution (pH 7.0-7.5) was sterile filtered with a hydrophobic PTFE membrane filter (pore size: 0.2 µm, Sartorius) before injection via a sterile 50 or 60 mm length 31 gauge sharp-tip steel cannula (Plastics One). The solution was delivered at a rate of 0.25 µl/min using a 0.1 ml glass-tight Hamilton syringe driven by a digital infusion pump (Harvard Apparatus). The injection volume per session was 4.5-5 µl of 10 mg/ml of THIP for monkey 1 (dPul in the left hemisphere) and 1.5 µl for monkey 2 (dPul in the right hemisphere). The injected volume was determined for each monkey separately in pilot sessions in which the monkey could perform the task without nystagmus and showed an ipsilesional spatial bias for target-target (free choice) stimuli as in prior reports (Wilke et al., 2010, 2013). The “free-choice” selection bias during target-target trials demonstrated a successful inactivation procedure.

Every experimental session started with a pre-injection testing period, followed by the injection, a 30-40 min waiting period, and the post-injection testing period. We conducted 7 inactivation sessions interleaved with 7 control (no actual injection) sessions for each monkey. In total, M1 performed 15222 trials and M2 performed 10945 trials. The control sessions were performed with the same timing of events as the inactivation sessions.

### Behavioral paradigm

The monkeys were sitting in a dark room in a custom-made primate chair with the head restrained 30 cm away from a 27” LED display (60 Hz refresh rate, model HN274H, Acer Inc. USA). The gaze position of the right eye was monitored at 220 Hz using an MCU02 ViewPoint infrared eye tracker (Arrington Research Inc. USA). A MATLAB-based task controller (https://github.com/dagdpz/monkeypsych, MATLAB version R2012b, The MathWorks, Inc., USA) and the Psychophysics Toolbox (Brainard, 1997) were used to control stimulus presentation.

#### Color discrimination task

Two monkeys performed a color discrimination task (**Figure 1B**) where the perceptual difficulty was determined by the color similarity of the target (T, red) vs. distractor (D, easy – yellow, difficult – orange). Target or distractor was either presented alone or with a second stimulus (distractor or target) in the opposite hemifield, determining the level of spatial competition. Monkeys had to saccade to the target or continue fixating when only distractor(s) were presented. Each trial started with the presentation of a red fixation spot. The monkey initiated each trial by acquiring eye fixation by entering the 5° radial window around the fixation spot within 500 ms after the onset of the fixation spot. After maintaining fixation for 500-900 ms, the fixation spot turned gray, and one or two peripheral dots simultaneously appeared (go-signal). Red dots represented targets, whereas yellow and orange dots represented distractors. In conditions with a single peripheral stimulus, either one target or one distractor was presented in the left or the right hemifield. In conditions with two peripheral stimuli, the monkey was shown two dots in opposite hemifields. In double-target trials, two equally-rewarded targets were presented, and the monkey could choose either as a saccade target. In double-distractor trials, two distractors were shown, which had to be ignored by maintaining central fixation. In target-distractor trials, a target was presented with a distractor in the opposite hemifield. The monkey was required to make a saccade towards the target while ignoring the distractor. The monkey had to choose within 500 ms (target acquisition epoch). As soon as the eye position entered the 5° radial window around one of the stimuli, the stimulus was selected, and the monkey was not allowed to reverse his decision. The chosen stimulus, either the selected peripheral dot for saccade responses or the fixation spot for maintaining eye fixation, turned bright to confirm the monkey’s selection. After fixating the selected stimulus for another 500 ms (target hold epoch), correct responses were followed by the reward tone, a fluid reward, and an inter-trial interval (ITI) of 2000 ms. Incorrect answers were followed by the error tone, no liquid reward, and an ITI of 2000 ms.

All stimuli were matched in luminance (dim stimuli: 11 cd/m^2^, bright stimuli: 35 cd/m^2^) and size (1° diameter). Targets and distractors were displayed at one of three locations per hemifield (six locations in total) with an eccentricity of 20° of visual angle. Stimulus locations were arranged concentrically around the fixation spot at 0° (mid-left), 20° (up left), 160° (upright), 180° (mid-right), 200° (down right), and 340° (down left). They were presented on a horizontal or a diagonal axis in conditions with two peripheral stimuli. All experimental conditions (stimulus type, difficulty level, spatial location) were pseudorandomized. Trials aborted before the monkey selected a stimulus returned to the pool of trials from which the next trial was chosen randomly.

#### Distractor color determination

After initial training of the task with an easy yellow distractor, we determined the difficult distractor color (orange) for the experiment based on the results of a psychophysical assessment (six sessions in each monkey). The goal of the assessment was to determine a distractor color that could be correctly discriminated from the target with 70 - 80% accuracy. To this end, the monkeys performed a color discrimination test with five distractor colors of different perceptual difficulty ranging from yellow (easy, RGB [60 60 0]) to red-orange (difficult, M1: [128 11 0]; M2: [128 23 0]). All trial conditions were presented in a pseudorandomized order. The perceptual difficulty was defined as the RGB value ratio between green (G) and red (R). A stimulus with a G/R ratio of 1 is a yellow distractor (60 60 0) which is perceptually very different from the target color ([128 0 0]) with a G/R ratio of 0. We chose an orange color as the difficult distractor color (M1: G/R ratio = 0.09, M2: G/R ratio = 0.18, see Supplementary Information, **Suppl. Figure S1**). The corresponding accuracy values were fitted by the cumulative normal function using Palamedes toolbox (Prins and Kingdom, 2009) in MATLAB 2019b (The MathWorks, Inc. USA).

#### Equalizing spatial choice behavior

In the beginning of each task training session, the left-right choice behavior in target-target trials was approximately equalized by shifting the entire stimulus array (both central and peripheral stimuli) with respect to the body midline, if a substantial hemifield bias was present (Dominguez-Vargas et al., 2017). In all subsequent experimental sessions, the stimulus array was adjusted by the same amount (0° from the midline for M1 and by 5° for M2).

## Data analysis

We analyzed the behavioral data from two monkeys for the “post-injection” testing period, with seven injections and seven control sessions for each monkey. Data analysis were performed using MATLAB R2015a. The analysis focuses on the saccade latencies, accuracy, and Signal Detection Theory variables to evaluate if there is a significant statistical difference between control and inactivation sessions. Accuracy was defined as the proportion of correct and rewarded trials among all trials in a specific condition (e.g. correct hits and rejections for all single stimuli trials, regardless of the hemifield). For calculating the saccade latencies, we used all completed saccades to the target.

### Saccade definition

All eye movements with a minimum velocity of 200 °/s and a minimum duration of 30 ms were considered saccades. To detect a saccade, the instantaneous saccade velocity was calculated sample by sample as the square root of the sum of squared interpolated (220 Hz to 1 kHz) and smoothed (12 ms moving average rectangular window) horizontal and vertical eye position traces, and smoothed again (12 ms moving average rectangular window). Saccade onset was defined as the first eye position change after the go-cue that exceeded a velocity threshold of 200°/s. Saccade end was defined as the first point in time when eye velocity dropped below 50°/s after saccade onset.

### Signal Detection Theory

To address our two alternative hypotheses, whether the contralesional visuospatial deficit is a consequence of a contralesional perceptual discrimination deficit or spatial selection bias, we used the Signal Detection Theory to assess changes in perceptual sensitivity index (d-prime) and response criterion after unilateral reversible dorsal pulvinar inactivation. D-prime measures how well the monkeys discriminate targets from distractors (Eq. 1); z represents z-score calculated using normal inverse cumulative distribution function (*norminv* function in MATLAB). The response criterion indicates the tendency to select a stimulus in a specific hemifield regardless if it is a target or distractor (Eq. 2).

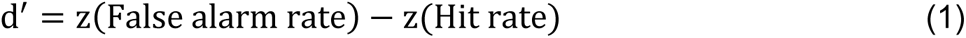

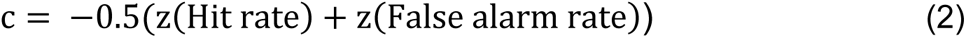

The data were analyzed separately for each monkey (M1 and M2), for each difficulty level (yellow and orange distractor), and stimulus type (single, double same, and double different stimuli). To compare the effect of inactivation per hemifield, we calculated the signal detection theory variables separately for the contralesional and ipsilesional hemifield relative to the side of inactivation (see the details in the Supplementary Information, **Suppl. Figures S2-S4**). An increase of the criterion signifies decreased selection of the contralateral stimulus (“less contra”) and vice versa.

### Statistical analysis

The statistical analysis was performed using R (version 4.1.2, R Core Team, 2022) and MATLAB 2014b. First, to assess whether the accuracy differs between the three stimulus types (single stimuli, double same, and double different stimuli) for each difficulty level in the control sessions due intended consequence of our task design, we conducted a mixed ANOVA followed by post-hoc tests to determine whether the three stimulus types differed significantly (corrected for multiple comparisons using Bonferroni correction).

The main aim of the study was to investigate the effects of dorsal pulvinar inactivation compared to control (no perturbation) sessions on different dependent variables. Accuracy was analyzed with three-way mixed ANOVAs: Difficulty level (easy, difficult) × Stimulus type (single, double same, double different) × Perturbation (inactivation, control). The d-prime and criterion were analyzed with four-way mixed ANOVAs: Difficulty level (easy, difficult) × Stimulus type (single, double same, double different) × Perturbation (inactivation, control) × Hemifield (contra, ipsi). Although the four-factor mixed ANOVA includes all possible interactions, it cannot directly answer our research question: whether dPul inactivation affects the criterion or d-prime, differently for the three stimulus types and the two perceptual difficulty levels. To assess whether there was a statistical difference between the inactivation sessions and control sessions in the d-prime and criterion, we conducted independent sample t-tests separately for the stimulus position (contralesional and ipsilesional hemifield), the stimulus type (single, double same, and double different), and the difficulty level (difficult and easy). Due to a small sample size, we also calculated non-parametric tests (Wilcoxon rank sum test), leading to comparable results (**Suppl. Table S8**).

### Simulations

We numerically simulated different scenarios of stimulus selection corresponding to the two alternative hypotheses (response bias and perceptual sensitivity deficit). These simulations aimed to visualize the effects of unilateral inactivation on selection behavior and resulting STD variables for each scenario, and to compare the changes derived from the predictions of each hypothesis with the data. One group of scenarios represents the response criterion hypothesis, where we expect a decrease both in contralesional hit rate and false alarm rate after the inactivation, resulting in a shift of criterion away from the contralesional hemifield – i.e. towards “less contra”. The other group of scenarios represents the perceptual discrimination hypothesis, where we expect a decrease in the contralesional hit rate and an increase in false alarm rate, resulting in decreased contralesional d-prime. In brief, the proportions of hits, misses, correct rejections, and false alarms were defined for the control condition (no inactivation), approximately based on the actual monkey performance. A specific bias or perceptual deficit on these proportions was introduced to estimate resulting hits, misses, correct rejections, and false alarms in the “inactivated” condition. The resulting criterion and d-prime values were calculated for the control and the inactivated conditions.

For each scenario, the simulated data consisted of 200 trials (100 target trials and 100 distractor trials) separated into four different outcomes: hits, misses, false alarms and correction rejections. The proportion of selection for one specific stimulus type and one difficulty level is set in each scenario. An example of a specific selection pattern is illustrated for the scenario “single stimulus – difficult distractor – response criterion hypothesis” (**Table 3**). Before inactivation, the (contralateral) selection pattern is: hits: 0.7 fraction of target selection (correct, 70 trials), misses: 0.3 fraction staying on the central fixation spot when a target is presented (incorrect, 30 trials); correct rejections 0.6 fraction staying when a difficult distractor is presented (correct, 60 trials) and false alarms: 0.4 fraction selecting the distractor (incorrect, 40 trials). The hit rate is 0.7, and the false alarm rate is 0.4 resulting in a criterion of −0.14 and a d-prime of 0.77. According to the prediction of response bias hypothesis, after the inactivation the monkey should select less often stimuli presented in the contralesional hemifield regardless whether a target or a distractor is shown. We therefore expect a decrease of hits and false alarms, and an increase of misses and correct rejections, resulting in a decrease of the contralesional hit rate and false alarm rate. In this example, we chose an inactivation-induced decrease in contralesional selection by 0.2 (20 trials) for the hit and false alarm rates, resulting in shift of the criterion towards “less contra” (increase of the criterion). Notably, the d-prime is also changing slightly. To visualize how the hit rate, the false alarm rate, the criterion and the d-prime are related, we visualized for each combination of the false alarm rate and the hit rate the resulting values of criterion and d-prime. All simulations were done in MATLAB. The code is publically available at https://github.com/dagdpz/perceptual-dis.

**Table 3.**
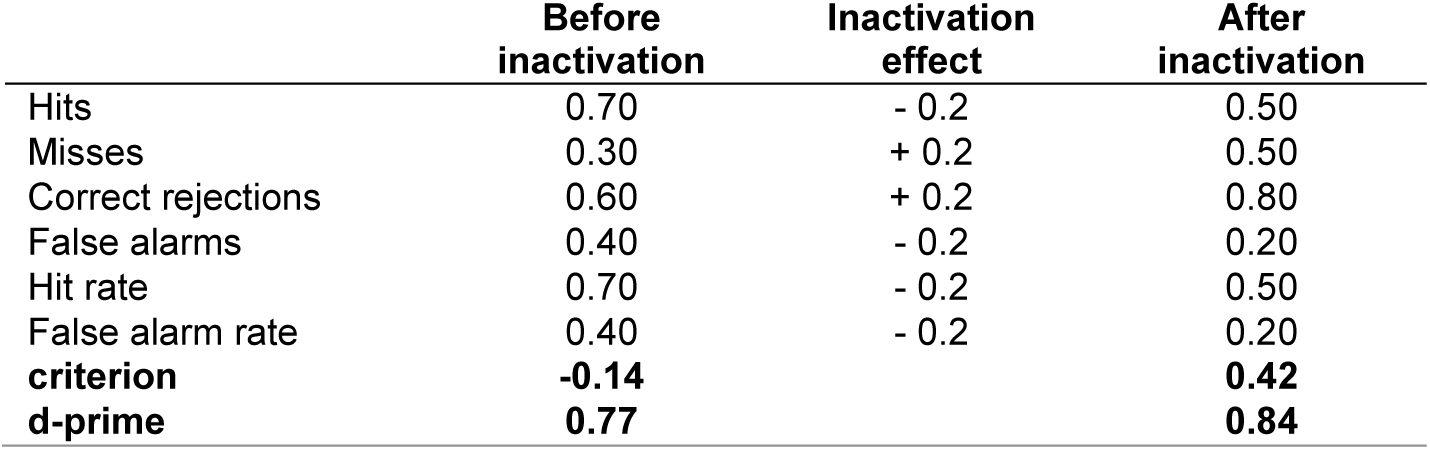
An example selection pattern for the simulated hypothetical scenario “single stimulus – difficult distractor – response bias hypothesis”. This example is illustrated in **Figure 4A**.

|  | **Before**<br>**inactivation** | **Inactivation**<br>**effect** | **After**<br>**inactivation** |
| --- | --- | --- | --- |
| Hits | 0.70 | - 0.2 | 0.50 |
| Misses | 0.30 | + 0.2 | 0.50 |
| Correct rejections | 0.60 | + 0.2 | 0.80 |
| False alarms | 0.40 | - 0.2 | 0.20 |
| Hit rate | 0.70 | - 0.2 | 0.50 |
| False alarm rate | 0.40 | - 0.2 | 0.20 |
| **criterion** | **-0.14** |  | **0.42** |
| **d-prime** | **0.77** |  | **0.84** |

## Authors contributions

Conceptualization: KK, MW, IK. Data curation: KK. Formal analysis: KK, IK. Funding acquisition: MW, IK. Project Administration: IK. Supervision: MW, IK. Visualization: KK, IK. Writing—Original draft preparation: KK, IK. Writing—Review & editing: KK, MW, IK.

## Acknowledgments

We thank Lydia Gibson and Uwe Zimmermann for developing the color discrimination task, Lukas Schneider for help with the monkeypsych experimental task software, and Daniela Lazzarini, Sina Plümer, Klaus Heisig, and Dirk Prüße for technical support. We also thank Stefan Treue, Alexander Gail, the Decision and Awareness Group, the Sensorimotor Group, and the Cognitive Neuroscience Laboratory for helpful discussions. Supported by the Hermann and Lilly Schilling Foundation, German Research Foundation (DFG) grants WI 4046/1-1 and Research Unit GA1475-B4, KA 3726/2-1, CNMPB Primate Platform, and core funding from the Cognitive Neuroscience Laboratory.

## Data and code availability statement

The datasets generated and analyzed for the current study, and the corresponding code, are available from the corresponding author on reasonable request.

---

**Supplementary Figure S1.**
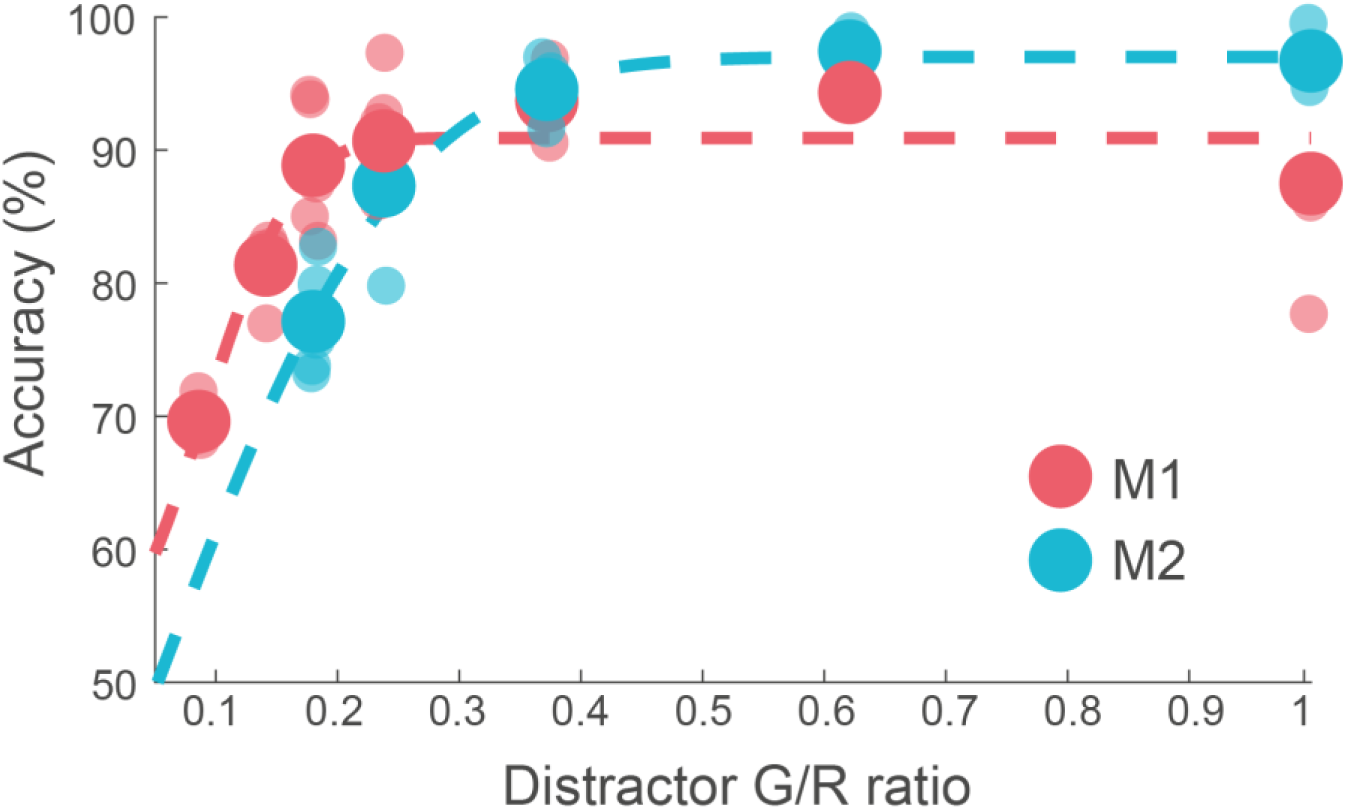
Accuracy per distractor color. The monkeys performed a color discrimination paradigm with five distractor colors of different perceptual difficulty ranging from yellow (easy, G/R ratio: 1, RGB [60 60 0]) to red-orange (difficult, G/R ratio M1: 0.09, M2: 0.18, [M1: 128 11 0; M2: 128 23 0]). We calculated how accurate the target was discriminated from a distractor in the opposite hemifield. The large dots display the mean accuracy across sessions for the different applied G/R ratios separated for each monkey. To these accuracy values, the cumulative normal function was fitted. The small transparent dots display the accuracy per session. The goal of the assessment was to determine a distractor color that could be correctly discriminated from the target with 70 - 80% accuracy, for the difficult perceptual condition.

**Supplementary Figure S2.**
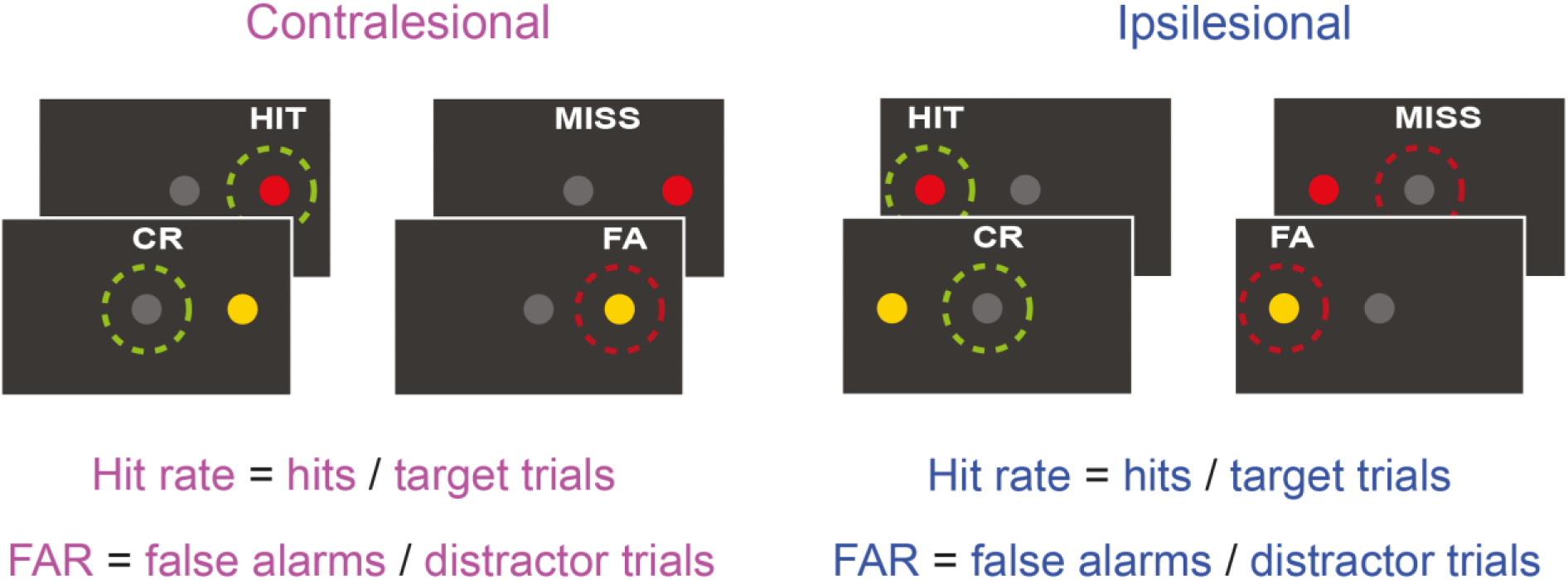
Calculation of the signal detection theory variables for single stimuli related to the results in Figure 4. Here we describe, firstly, how trials were classified in relation to the monkey’s responses and secondly, the calculations. Hits are trials where a saccade was made to a target (correct response, green dashed circle). Misses are trials where the monkey fixated the dot in the middle of the screen while a single target was displayed (incorrect response, dark red dashed circle). Correct rejections are trials where the monkey fixated the dot in the middle of the screen when a single distractor was displayed in the periphery (correct response, green dashed circle). False alarms are trials where the monkey made a saccade to the distractor (incorrect response, dark red dashed circle). We calculated the hit rate and false alarm rate (FAR) according to Hit rate = Hits / contralesional target trials and False alarm rate = False alarms / contralesional distractor trials. We used the standard calculations for the d-prime (d’ = z (Hit) –z(FAR)) and criterion (c = −0.5*(z(Hit) + z(FAR))). All variables were calculated separately for stimuli presented in the ipsilesional and contralesional hemifield to compare the changes in d-prime and criterion for each hemifield.

**Supplementary Figure S3.**
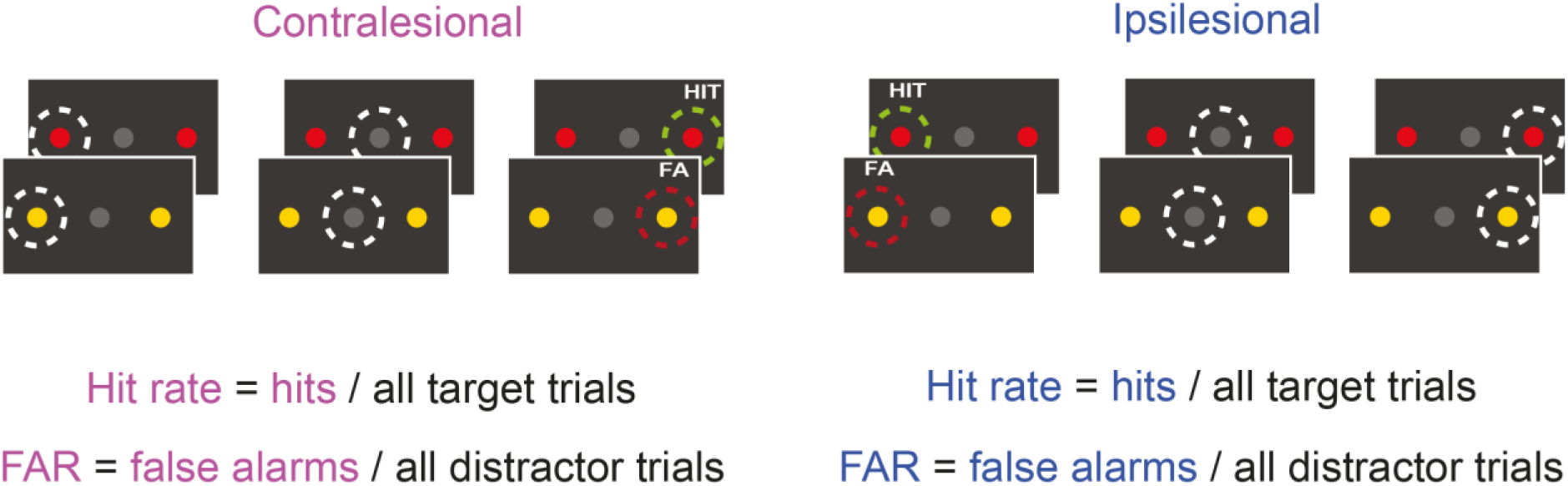
Calculation of the signal detection theory variables for double same stimuli related to the results in Figure 5. Notations are the same as in **Suppl. Figure 2**. All variables were calculated separately for stimuli presented in the ipsilesional or contralesional hemifield, which allows us to compare the changes in d-prime and criterion for each hemifield. In the following, the examples are given for the contralesional hemifield. Contralesional hits are trials where a saccade was made to the contralesional target when a target was presented in each hemifield. Contralesional false alarms are trials where a saccade was made to the contralesional distractor when a distractor was presented in each hemifield. Hit rate is computed as all contralesional hits divided by all double same target trials. False alarm rate is computed as all contralesional false alarms divided by all double same distractor trials. We used the standard calculations for the d-prime (d’ = z(Hit) –z(FAR)) and criterion (c = - 0.5*(z(Hit) + z(FAR))).

**Supplementary Figure S4.**
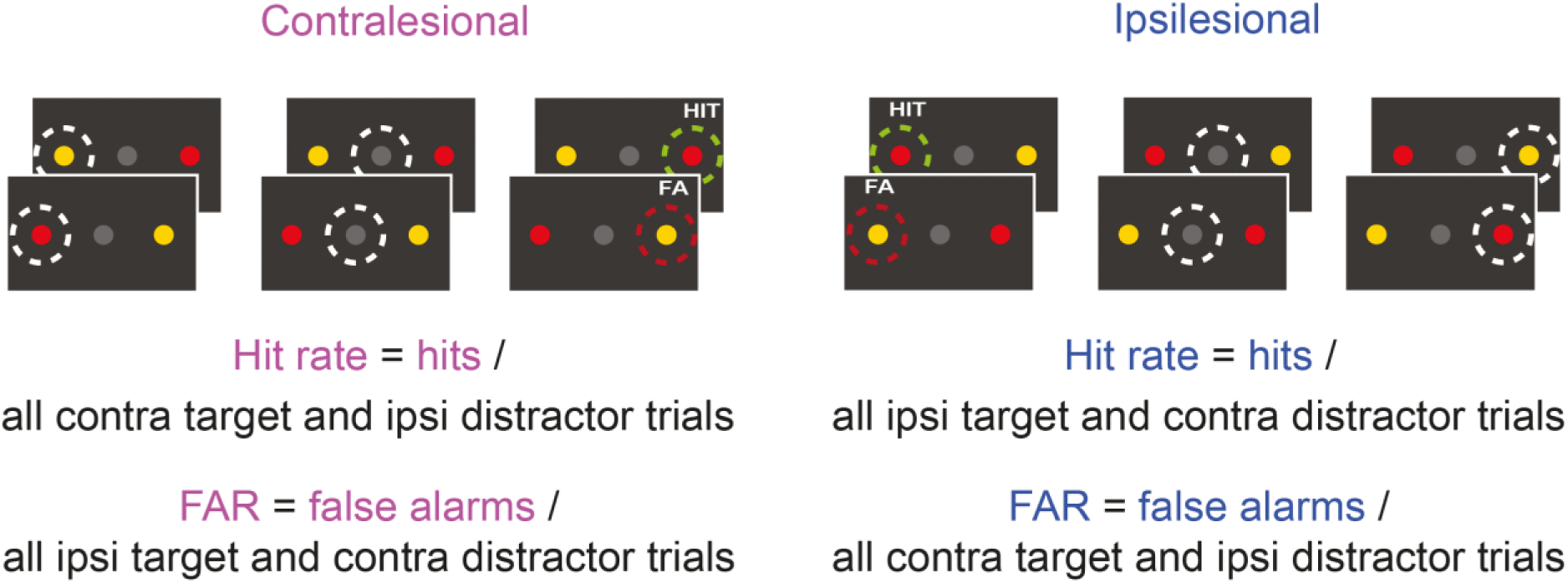
Calculation of the signal detection theory variables for double different stimuli related to the results in Figure 6. Notations are the same as in **Suppl. Figure 2**. All variables were calculated separately for stimuli presented in the ipsilesional or contralesional hemifield, which allows us to compare the changes in d-prime and criterion for each hemifield. In the following, examples are given for the ipsilesional hemifield. Ipsilesional hits are trials where a saccade was made to the ipsilesional target. Ipsilesional false alarms are trials where a saccade was made to the ipsilesional distractor. The hit rate is computed as all ipsilesional hits divided by all double different trials where a target was presented in the ipsilesional hemifield (including all response options, i.e. fixation and saccades to either ipsi- or contralesional stimulus). Likewise, false alarm rate (FAR) is computed as all ipsilesional false alarms divided by all double different trials where a distractor was presented in the ipsilesional hemifield. We used the standard calculations for the d-prime (d’ = z(Hit) –z(FAR)) and criterion (c = −0.5*(z(Hit) + z(FAR)).

**Supplementary Figure S5.**
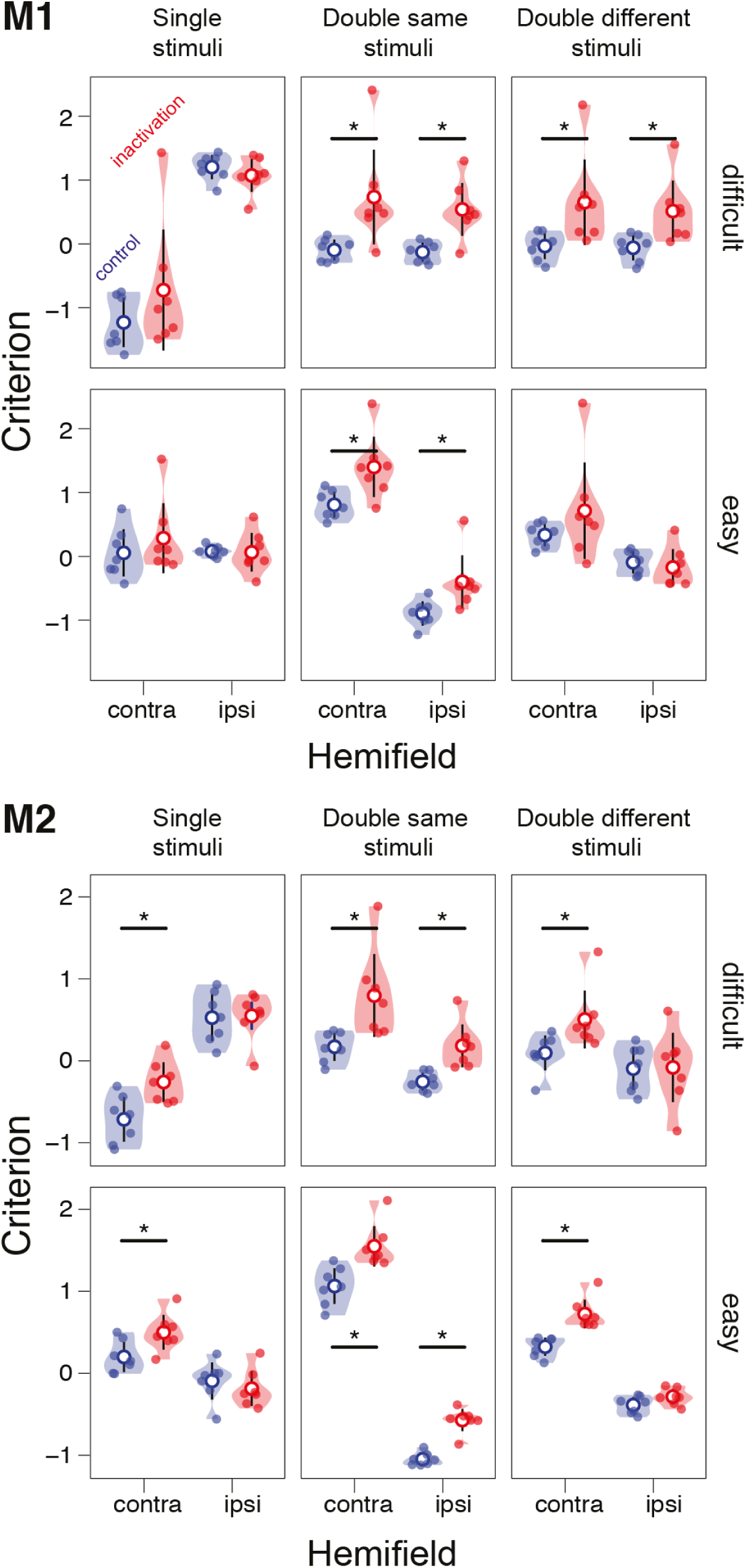
Summary of the inactivation results for the criterion. The violin plots display the distribution of the computed values for the criterion for the two difficulty levels (difficult and easy), stimulus types (single / double same / double different stimuli), and hemifield (contra-/ipsilesional) for control (blue) and inactivation (red) sessions. The mean is displayed with an empty circle and 95% confidence intervals are shown. Each small colored dot represents a session. The stars display the significance of the t-test at the p-value < 0.05.

**Supplementary Figure S6.**
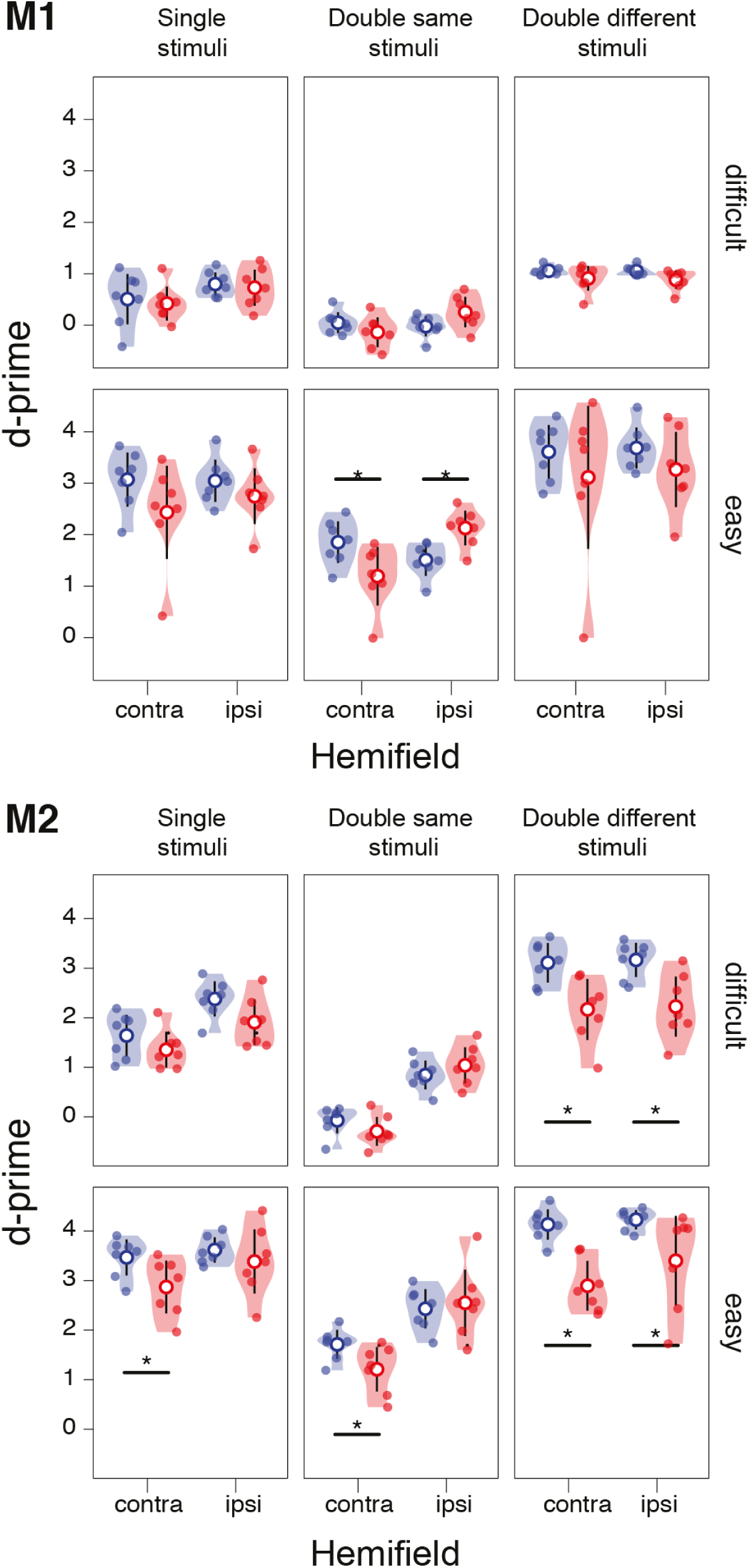
Summary of the inactivation results for the d-prime. The violin plots display the distribution of the computed values for the criterion for the two difficulty levels (difficult and easy), stimulus types (single / double same / double different stimuli), and hemifield (contra-/ipsilesional) for control (blue) and inactivation (red) sessions. The mean is displayed with an empty circle and 95% confidence intervals are shown. Each small colored dot represents a session. The stars display the significance of the t-test at the p-value < 0.05.

**Supplementary Table S1.** Three-way mixed ANOVA on saccade latency, with within-factors “Stimulus Type” (single / double same / double different) and “Hemifield” (contralesional/ipsilesional) and between-factor “Perturbation” (control/inactivation sessions); ges - generalized eta squared. Significant effects are shown in bold font.

| Factor | DFn | DFd | F | p-value | p<.05 | ges | Monkey |
| --- | --- | --- | --- | --- | --- | --- | --- |
| **Perturbation** | **1** | **11** | **5.96** | **0.033** | **\*** | **0.24** | M1 |
| **Hemifield** | **1** | **11** | **20.08** | **0.001** | **\*** | **0.18** |  |
| **Stimulus Type** | **2** | **22** | **25.29** | **< .001** | **\*** | **0.20** |  |
| **Perturbation × Hemifield** | **1** | **11** | **21.81** | **0.001** | **\*** | **0.20** |  |
| **Perturbation × Stimulus Type** | **2** | **22** | **5.25** | **0.014** | **\*** | **0.05** |  |
| Hemifield × Stimulus Type | 2 | 22 | 1.12 | 0.344 |  | 0.02 |  |
| **Perturbation × Hemifield × Stimulus Type** | **2** | **22** | **4.64** | **0.021** | **\*** | **0.07** |  |
| **Perturbation** | **1** | **12** | **7.48** | **0.018** | **\*** | **0.26** | M2 |
| **Hemifield** | **1** | **12** | **6.72** | **0.024** | **\*** | **0.15** |  |
| **Stimulus Type** | **2** | **24** | **14.19** | **< .001** | **\*** | **0.08** |  |
| Perturbation × Hemifield | 1 | 12 | 3.47 | 0.087 |  | 0.08 |  |
| Perturbation × Stimulus Type | 2 | 24 | 1.27 | 0.299 |  | 0.01 |  |
| **Hemifield × Stimulus Type** | **2** | **24** | **11.44** | **< .001** | **\*** | **0.05** |  |
| **Perturbation × Hemifield × Stimulus Type** | **2** | **24** | **3.74** | **0.039** | **\*** | **0.02** |  |

**Supplementary Table S2.**
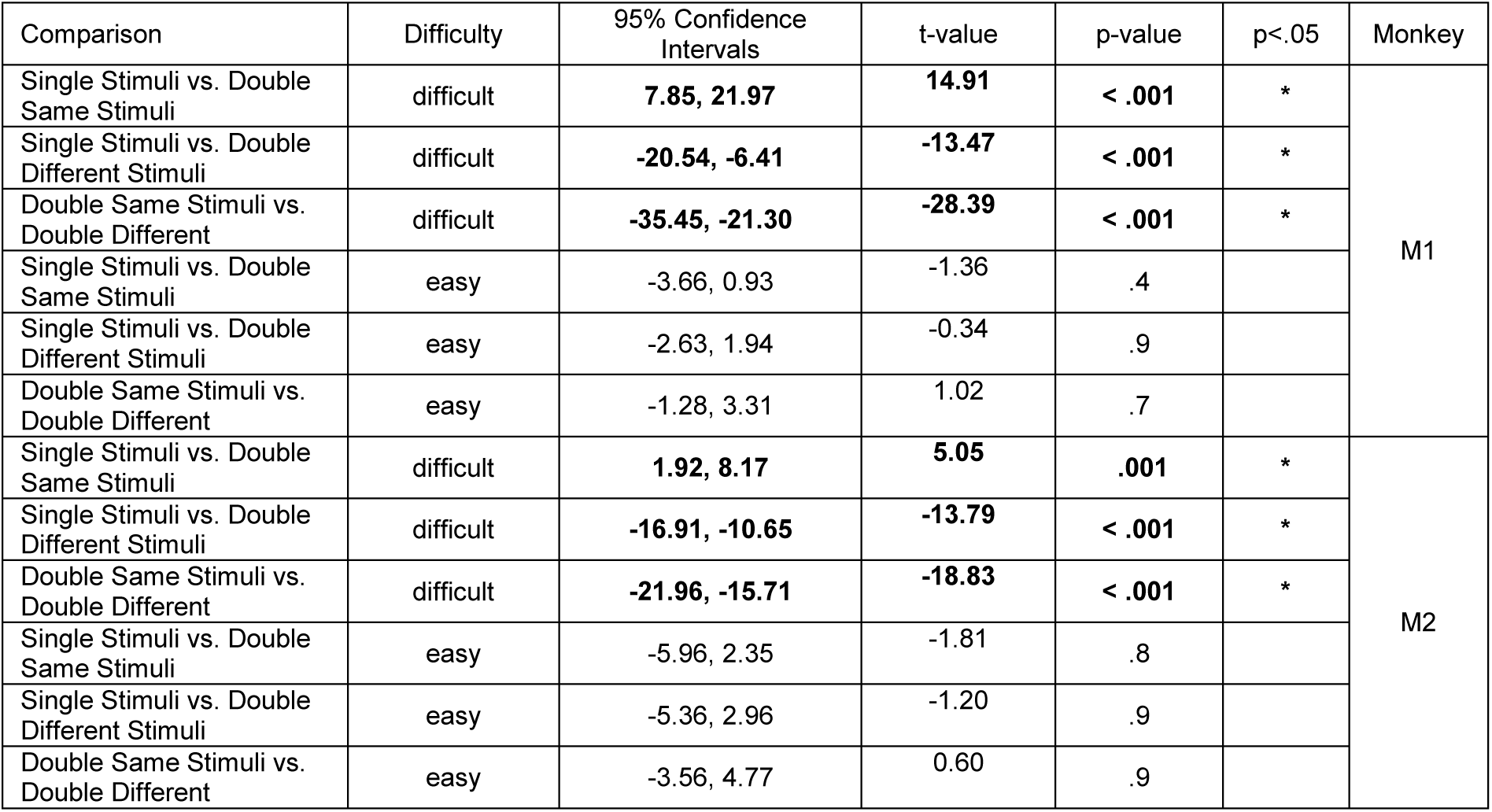
Pairwise t-tests comparing the accuracy in the different stimulus type conditions for the control sessions, separately within each perceptual difficulty (difficult/easy discrimination). Significant effects are shown in bold font.

**Supplementary Table S3.** Three-way mixed ANOVA on accuracy, with within-factors “Stimulus Type” (single / double same / double different and “Difficulty” (difficult/easy discrimination) and between-factor “Perturbation” (control/inactivation sessions); ges - generalized eta squared. Significant effects are shown in bold font.

| Factor | DFn | DFd | F | p-value | p<.05 | ges | Monkey |
| --- | --- | --- | --- | --- | --- | --- | --- |
| Perturbation | 1 | 12 | 1.46 | 0.250 |  | 0.06 | M1 |
| Difficulty | 1 | 12 | 349.71 | < .001 | * | 0.87 |  |
| Stimulus Type | 2 | 24 | 6.09 | .007 | * | 0.11 |  |
| Perturbation × Difficulty | 1 | 12 | 1.36 | .275 |  | 0.03 |  |
| Perturbation × Stimulus Type | 2 | 24 | 3.17 | .060 |  | 0.06 |  |
| Difficulty × Stimulus Type | 2 | 24 | 124.01 | < .001 | * | 0.29 |  |
| Perturbation × Difficulty × Stimulus Type | 2 | 24 | 1.88 | .175 |  | 0.01 |  |
| Perturbation | 1 | 12 | 9.42 | .010 | * | 0.22 | M2 |
| Difficulty | 1 | 12 | 421.89 | < .001 | * | 0.72 |  |
| Stimulus Type | 2 | 24 | 8.12 | .002 | * | 0.24 |  |
| Perturbation × Difficulty | 1 | 12 | 0.07 | .798 |  | 0.0004 |  |
| Perturbation × Stimulus Type | 2 | 24 | 6.40 | .006 | * | 0.20 |  |
| Difficulty × Stimulus Type | 2 | 24 | 104.36 | < .001 | * | 0.48 |  |
| Perturbation × Difficulty × Stimulus Type | 2 | 24 | 1.12 | .342 |  | 0.01 |  |

**Supplementary Table S4.**
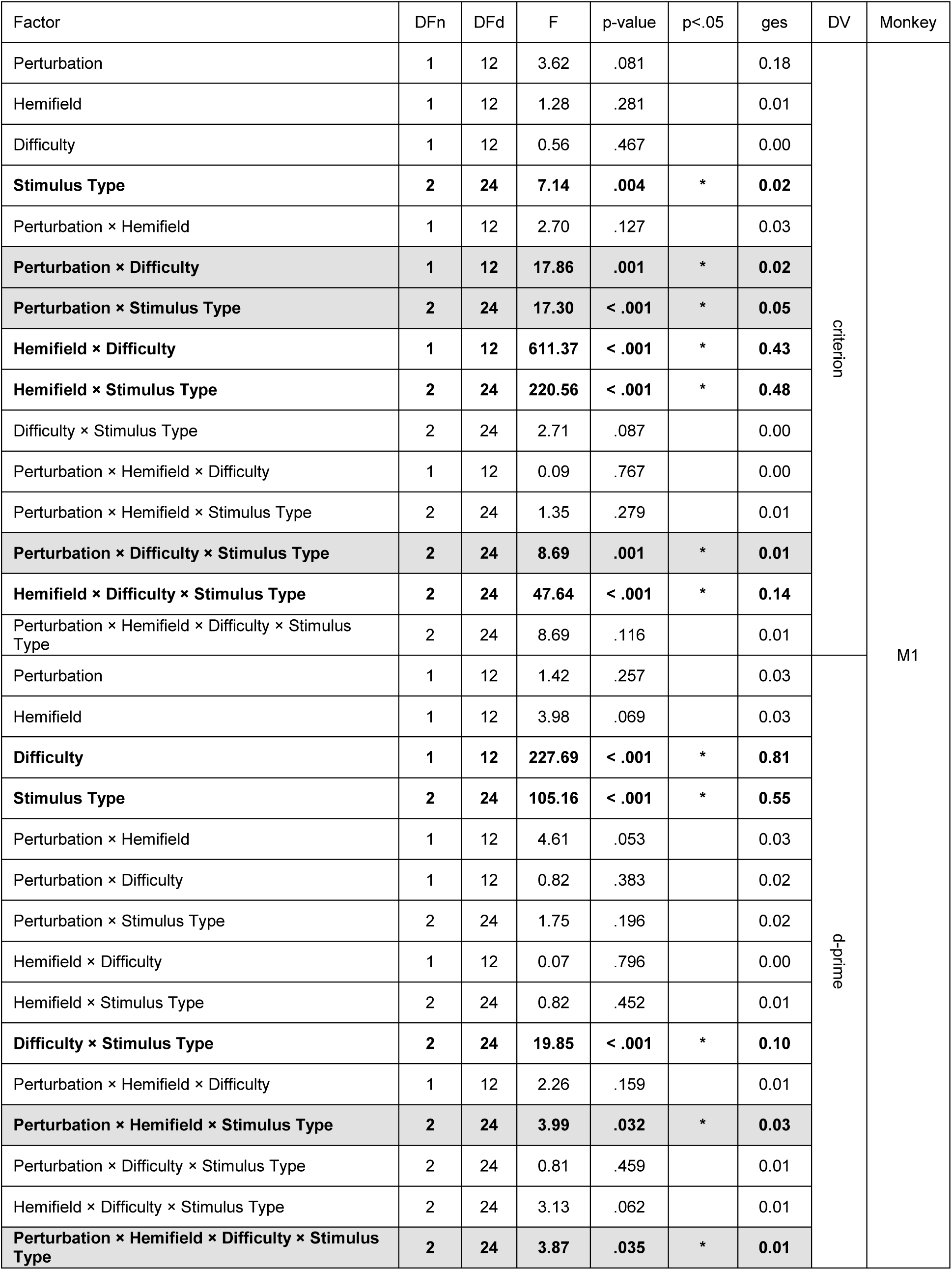
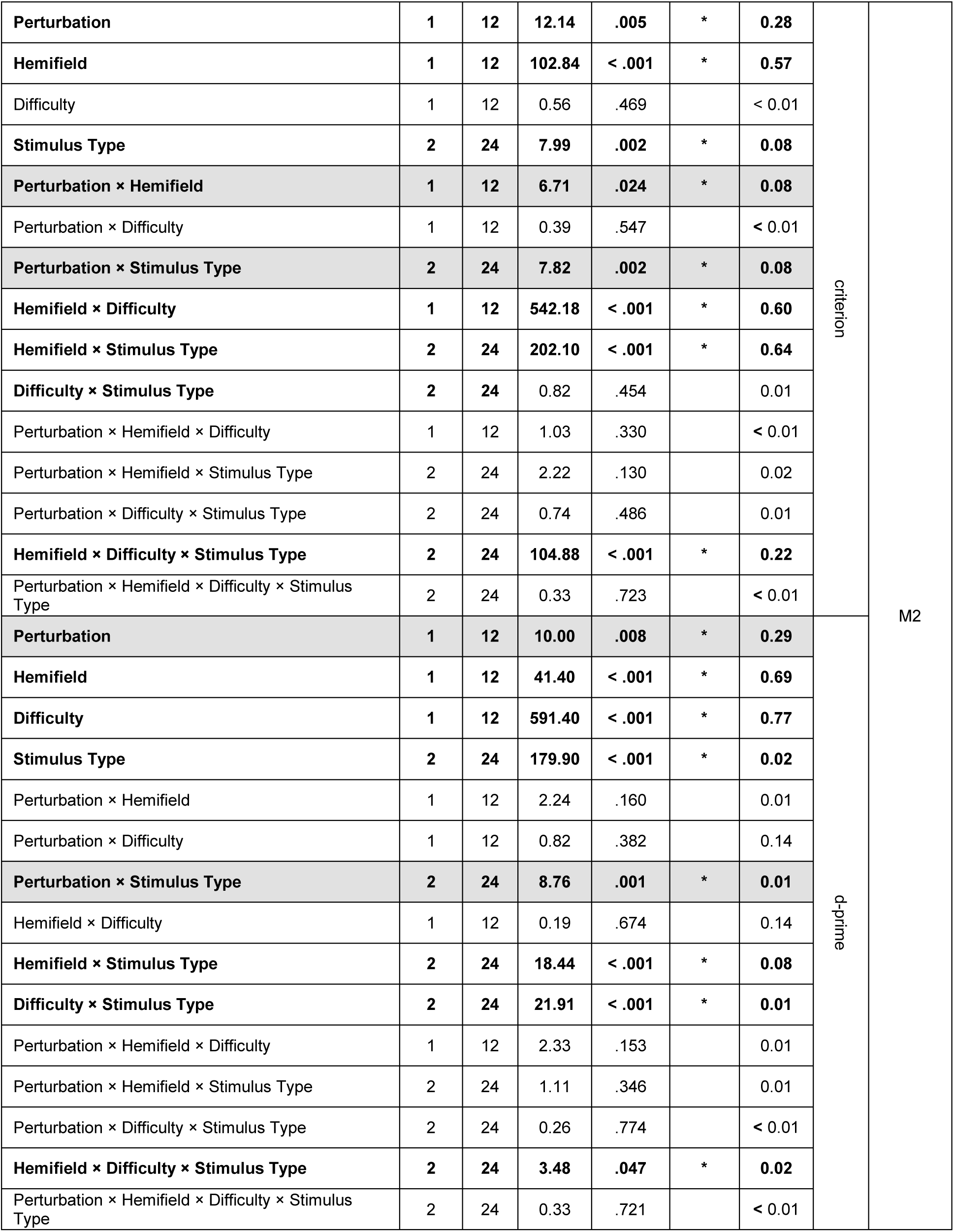
. Four-way mixed ANOVA on d-prime and criterion with within-factor “Stimulus type” (single / double same / double different), “Difficulty” (easy / difficult), and “Hemifield” (contralesional/ ipsilesional) and between-factor “Perturbation” (control / inactivation sessions), separately for each stimulus type, for the difficult discrimination; ges - generalized eta squared. Significant effects are shown in bold font, effects involving perturbation factor are highlighted with gray background.

| Factor | DFn | DFd | F | p-value | p<.05 | ges | DV | Monkey |
| --- | --- | --- | --- | --- | --- | --- | --- | --- |
| Perturbation | 1 | 12 | 3.62 | .081 |  | 0.18 | criterion | M1 |
| Hemifield | 1 | 12 | 1.28 | .281 |  | 0.01 |  |  |
| Difficulty | 1 | 12 | 0.56 | .467 |  | 0.00 |  |  |
| Stimulus Type | 2 | 24 | 7.14 | .004 | * | 0.02 |  |  |
| Perturbation × Hemifield | 1 | 12 | 2.70 | .127 |  | 0.03 |  |  |
| Perturbation × Difficulty | 1 | 12 | 17.86 | .001 | * | 0.02 |  |  |
| Perturbation × Stimulus Type | 2 | 24 | 17.30 | < .001 | * | 0.05 |  |  |
| Hemifield × Difficulty | 1 | 12 | 611.37 | < .001 | * | 0.43 |  |  |
| Hemifield × Stimulus Type | 2 | 24 | 220.56 | < .001 | * | 0.48 |  |  |
| Difficulty × Stimulus Type | 2 | 24 | 2.71 | .087 |  | 0.00 |  |  |
| Perturbation × Hemifield × Difficulty | 1 | 12 | 0.09 | .767 |  | 0.00 |  |  |
| Perturbation × Hemifield × Stimulus Type | 2 | 24 | 1.35 | .279 |  | 0.01 |  |  |
| Perturbation × Difficulty × Stimulus Type | 2 | 24 | 8.69 | .001 | * | 0.01 |  |  |
| Hemifield × Difficulty × Stimulus Type | 2 | 24 | 47.64 | < .001 | * | 0.14 |  |  |
| Perturbation × Hemifield × Difficulty × Stimulus Type | 2 | 24 | 8.69 | .116 |  | 0.01 |  |  |
| Perturbation | 1 | 12 | 1.42 | .257 |  | 0.03 | d-prime |  |
| Hemifield | 1 | 12 | 3.98 | .069 |  | 0.03 |  |  |
| Difficulty | 1 | 12 | 227.69 | < .001 | * | 0.81 |  |  |
| Stimulus Type | 2 | 24 | 105.16 | < .001 | * | 0.55 |  |  |
| Perturbation × Hemifield | 1 | 12 | 4.61 | .053 |  | 0.03 |  |  |
| Perturbation × Difficulty | 1 | 12 | 0.82 | .383 |  | 0.02 |  |  |
| Perturbation × Stimulus Type | 2 | 24 | 1.75 | .196 |  | 0.02 |  |  |
| Hemifield × Difficulty | 1 | 12 | 0.07 | .796 |  | 0.00 |  |  |
| Hemifield × Stimulus Type | 2 | 24 | 0.82 | .452 |  | 0.01 |  |  |
| Difficulty × Stimulus Type | 2 | 24 | 19.85 | < .001 | * | 0.10 |  |  |
| Perturbation × Hemifield × Difficulty | 1 | 12 | 2.26 | .159 |  | 0.01 |  |  |
| Perturbation × Hemifield × Stimulus Type | 2 | 24 | 3.99 | .032 | * | 0.03 |  |  |
| Perturbation × Difficulty × Stimulus Type | 2 | 24 | 0.81 | .459 |  | 0.01 |  |  |
| Hemifield × Difficulty × Stimulus Type | 2 | 24 | 3.13 | .062 |  | 0.01 |  |  |
| Perturbation × Hemifield × Difficulty × Stimulus Type | 2 | 24 | 3.87 | .035 | * | 0.01 |  |  |
| **Perturbation** | **1** | **12** | **12.14** | **.005** | **\*** | **0.28** | criterion | M2 |
| **Hemifield** | **1** | **12** | **102.84** | **< .001** | **\*** | **0.57** |  |  |
| Difficulty | 1 | 12 | 0.56 | .469 |  | < 0.01 |  |  |
| **Stimulus Type** | **2** | **24** | **7.99** | **.002** | **\*** | **0.08** |  |  |
| **Perturbation × Hemifield** | **1** | **12** | **6.71** | **.024** | **\*** | **0.08** |  |  |
| Perturbation × Difficulty | 1 | 12 | 0.39 | .547 |  | < 0.01 |  |  |
| **Perturbation × Stimulus Type** | **2** | **24** | **7.82** | **.002** | **\*** | **0.08** |  |  |
| **Hemifield × Difficulty** | **1** | **12** | **542.18** | **< .001** | **\*** | **0.60** |  |  |
| **Hemifield × Stimulus Type** | **2** | **24** | **202.10** | **< .001** | **\*** | **0.64** |  |  |
| **Difficulty × Stimulus Type** | **2** | **24** | 0.82 | .454 |  | 0.01 |  |  |
| Perturbation × Hemifield × Difficulty | 1 | 12 | 1.03 | .330 |  | < 0.01 |  |  |
| Perturbation × Hemifield × Stimulus Type | 2 | 24 | 2.22 | .130 |  | 0.02 |  |  |
| Perturbation × Difficulty × Stimulus Type | 2 | 24 | 0.74 | .486 |  | 0.01 |  |  |
| **Hemifield × Difficulty × Stimulus Type** | **2** | **24** | **104.88** | **< .001** | **\*** | **0.22** |  |  |
| Perturbation × Hemifield × Difficulty × Stimulus Type | 2 | 24 | 0.33 | .723 |  | < 0.01 |  |  |
| **Perturbation** | **1** | **12** | **10.00** | **.008** | **\*** | **0.29** | d-prime |  |
| **Hemifield** | **1** | **12** | **41.40** | **< .001** | **\*** | **0.69** |  |  |
| **Difficulty** | **1** | **12** | **591.40** | **< .001** | **\*** | **0.77** |  |  |
| **Stimulus Type** | **2** | **24** | **179.90** | **< .001** | **\*** | **0.02** |  |  |
| Perturbation × Hemifield | 1 | 12 | 2.24 | .160 |  | 0.01 |  |  |
| Perturbation × Difficulty | 1 | 12 | 0.82 | .382 |  | 0.14 |  |  |
| **Perturbation × Stimulus Type** | **2** | **24** | **8.76** | **.001** | **\*** | **0.01** |  |  |
| Hemifield × Difficulty | 1 | 12 | 0.19 | .674 |  | 0.14 |  |  |
| **Hemifield × Stimulus Type** | **2** | **24** | **18.44** | **< .001** | **\*** | **0.08** |  |  |
| **Difficulty × Stimulus Type** | **2** | **24** | **21.91** | **< .001** | **\*** | **0.01** |  |  |
| Perturbation × Hemifield × Difficulty | 1 | 12 | 2.33 | .153 |  | 0.01 |  |  |
| Perturbation × Hemifield × Stimulus Type | 2 | 24 | 1.11 | .346 |  | 0.01 |  |  |
| Perturbation × Difficulty × Stimulus Type | 2 | 24 | 0.26 | .774 |  | < 0.01 |  |  |
| **Hemifield × Difficulty × Stimulus Type** | **2** | **24** | **3.48** | **.047** | **\*** | **0.02** |  |  |
| Perturbation × Hemifield × Difficulty × Stimulus Type | 2 | 24 | 0.33 | .721 |  | < 0.01 |  |  |

**Supplementary Table S5.** Two-way mixed ANOVA on d-prime and criterion with within-factor “Hemifield” (contralesional/ipsilesional) and between-factor “Perturbation” (control/inactivation sessions), separately for each stimulus type, **for the difficult discrimination**; ges - generalized eta squared. Significant effects are shown in bold font.

| Factor | DFn | DFd | F | p-value | p<.05 | ges | DV | Monkey | Stimulus type |
| --- | --- | --- | --- | --- | --- | --- | --- | --- | --- |
| Perturbation | 1 | 12 | 0.70 | .42 |  | 0.03 | criterion | M1 | Single |
| Hemifield | 1 | 12 | 98.70 | < .001 | * | 0.80 |  |  |  |
| Perturbation × Hemifield | 1 | 12 | 2.23 | .161 |  | 0.08 |  |  |  |
| Perturbation | 1 | 12 | 0.31 | .588 |  | 0.01 | d-prime |  |  |
| Hemifield | 1 | 12 | 3.77 | .076 |  | 0.15 |  |  |  |
| Perturbation × Hemifield | 1 | 12 | 0.003 | .96 |  | < .001 |  |  |  |
| Perturbation | 1 | 12 | 6.40 | .026 | * | 0.17 | criterion | M2 |  |
| Hemifield | 1 | 12 | 69.56 | < .001 | * | 0.78 |  |  |  |
| Perturbation × Hemifield | 1 | 12 | 3.09 | .104 |  | 0.14 |  |  |  |
| Perturbation | 1 | 12 | 3.46 | .087 |  | 0.18 | d-prime |  |  |
| Hemifield | 1 | 12 | 33.58 | < .001 | * | 0.39 |  |  |  |
| Perturbation × Hemifield | 1 | 12 | 0.68 | .425 |  | 0.013 |  |  |  |
| Perturbation | 1 | 12 | 9.65 | .009 | * | 0.42 | criterion | M1 | Double Same |
| Hemifield | 1 | 12 | 2.19 | .165 |  | 0.02 |  |  |  |
| Perturbation × Hemifield | 1 | 12 | 1.11 | .313 |  | 0.01 |  |  |  |
| Perturbation | 1 | 12 | 0.76 | .401 |  | 0.01 | d-prime |  |  |
| Hemifield | 1 | 12 | 1.42 | .256 |  | 0.09 |  |  |  |
| Perturbation × Hemifield | 1 | 12 | 2.89 | .115 |  | 0.17 |  |  |  |
| Perturbation | 1 | 12 | 10.43 | .007 | * | 0.44 | criterion | M2 |  |
| Hemifield | 1 | 12 | 82.07 | < .001 | * | 0.43 |  |  |  |
| Perturbation × Hemifield | 1 | 12 | 2.62 | .132 |  | 0.02 |  |  |  |
| Perturbation | 1 | 12 | 0.02 | .888 |  | 0.00 | d-prime |  |  |
| Hemifield | 1 | 12 | 55.80 | < .001 | * | 0.77 |  |  |  |
| Perturbation × Hemifield | 1 | 12 | 1.95 | .188 |  | 0.11 |  |  |  |
| Perturbation | 1 | 12 | 6.47 | .026 | * | 0.34 | criterion | M1 | Double Different |
| Hemifield | 1 | 12 | 4.04 | .067 |  | 0.01 |  |  |  |
| Perturbation × Hemifield | 1 | 12 | 1.86 | .198 |  | 0.00 |  |  |  |
| Perturbation | 1 | 12 | 2.91 | .114 |  | 0.19 | d-prime |  |  |
| Hemifield | 1 | 12 | 0.90 | .361 |  | 0.00 |  |  |  |
| Perturbation × Hemifield | 1 | 12 | 0.32 | .582 |  | 0.00 |  |  |  |
| Perturbation | 1 | 12 | 1.53 | .24 |  | 0.10 | criterion | M2 |  |
| Hemifield | 1 | 12 | 26.77 | < .001 | * | 0.27 |  |  |  |
| Perturbation × Hemifield | 1 | 12 | 6.88 | .022 | * | 0.09 |  |  |  |
| Perturbation | 1 | 12 | 10.73 | .007 | * | 0.46 | d-prime |  |  |
| Hemifield | 1 | 12 | 1.23 | .289 |  | < 0.01 |  |  |  |
| Perturbation × Hemifield | 1 | 12 | < 0.01 | .993 |  | < 0.01 |  |  |  |

**Supplementary Table S6.** Two-way mixed ANOVA on d-prime and criterion with within-factor “Hemifield” (contralesional/ipsilesional) and between-factor “Perturbation” (control/inactivation sessions), separately for each stimulus type, **for the easy discrimination**; ges - generalized eta squared. Significant effects are shown in bold font.

| Factor | DFn | DFd | F | p-value | p<.05 | ges | DV | Monkey | Stimulus type |
| --- | --- | --- | --- | --- | --- | --- | --- | --- | --- |
| Perturbation | 1 | 12 | 0.37 | .557 |  | 0.02 | criterion | M1 | Single |
| Hemifield | 1 | 12 | 0.74 | .405 |  | 0.02 |  |  |  |
| Perturbation × Hemifield | 1 | 12 | 1.14 | .306 |  | 0.03 |  |  |  |
| Perturbation | 1 | 12 | 2.09 | .174 |  | 0.12 | d-prime |  |  |
| Hemifield | 1 | 12 | 0.84 | .378 |  | 0.01 |  |  |  |
| Perturbation × Hemifield | 1 | 12 | 1.16 | .302 |  | 0.02 |  |  |  |
| Perturbation | 1 | 12 | 1.73 | .213 |  | 0.06 | criterion | M2 |  |
| **Hemifield** | **1** | **12** | **28.33** | **< .001** | **\*** | **0.58** |  |  |  |
| Perturbation × Hemifield | 1 | 12 | 4.59 | .053 |  | 0.18 |  |  |  |
| Perturbation | 1 | 12 | 3.50 | .086 |  | 0.16 | d-prime |  |  |
| Hemifield | 1 | 12 | 4.34 | .059 |  | 0.11 |  |  |  |
| Perturbation × Hemifield | 1 | 12 | 1.28 | .279 |  | 0.04 |  |  |  |
| **Perturbation** | **1** | **12** | **8.30** | **.014** | **\*** | **0.39** | criterion | M1 | Double Same |
| **Hemifield** | **1** | **12** | **832.64** | **< .001** | **\*** | **0.87** |  |  |  |
| Perturbation × Hemifield | 1 | 12 | 0.62 | .445 |  | 0.01 |  |  |  |
| Perturbation | 1 | 12 | 0.02 | .899 |  | 0.001 | d-prime |  |  |
| Hemifield | 1 | 12 | 2.56 | .135 |  | 0.11 |  |  |  |
| **Perturbation × Hemifield** | **1** | **12** | **11.99** | **.005** | **\*** | **0.37** |  |  |  |
| **Perturbation** | **1** | **12** | **31.20** | **< .001** | **\*** | **0.64** | criterion | M2 |  |
| **Hemifield** | **1** | **12** | **1252.30** | **< .001** | **\*** | **0.97** |  |  |  |
| Perturbation × Hemifield | 1 | 12 | 0.004 | .949 |  | < 0.01 |  |  |  |
| Perturbation | 1 | 12 | 2.05 | .178 |  | 0.04 | d-prime |  |  |
| **Hemifield** | **1** | **12** | **18.84** | **< .001** | **\*** | **0.54** |  |  |  |
| Perturbation × Hemifield | 1 | 12 | 1.72 | .215 |  | 0.09 |  |  |  |
| Perturbation | 1 | 12 | 0.49 | .498 |  | 0.03 | criterion | M1 | Double Different |
| **Hemifield** | **1** | **12** | **32.22** | **< .001** | **\*** | **0.38** |  |  |  |
| Perturbation × Hemifield | 1 | 12 | 3.93 | .071 |  | 0.07 |  |  |  |
| Perturbation | 1 | 12 | 0.98 | .342 |  | 0.07 | d-prime |  |  |
| Hemifield | 1 | 12 | 0.46 | .511 |  | < 0.01 |  |  |  |
| Perturbation × Hemifield | 1 | 12 | 0.05 | .823 |  | < 0.01 |  |  |  |
| **Perturbation** | **1** | **12** | **22.96** | **< .001** | **\*** | **0.51** | criterion | M2 |  |
| **Hemifield** | **1** | **12** | **318.79** | **< .001** | **\*** | **0.92** |  |  |  |
| **Perturbation × Hemifield** | **1** | **12** | **9.52** | **.009** | **\*** | **0.27** |  |  |  |
| **Perturbation** | **1** | **12** | **13.49** | **.003** | **\*** | **0.47** | d-prime |  |  |
| Hemifield | 1 | 12 | 4.24 | .062 |  | 0.07 |  |  |  |
| Perturbation × Hemifield | 1 | 12 | 1.94 | .189 |  | 0.03 |  |  |  |

**Supplementary Table S7.** Non-hemifield-specific bias (“stay” – central fixation option, vs. “go” – saccade to a peripheral stimulus) in control and inactivation sessions. The negative criterion indicated a “go” bias. The table shows the results of the two-sided t-test against zero.

| Monkey | Difficulty | Session type | Stimulus type | Criterion | t-value | p-value | Bias |
| --- | --- | --- | --- | --- | --- | --- | --- |
| M1 | difficult | control | single stimuli | -1.21 | -11.16 | < .001 | Go |
|  |  | inactivation |  | -0.89 | -4.73 | .003 | Go |
|  | easy | control |  | 0.01 | 0.13 | .903 | Neutral |
|  |  | inactivation |  | 0.10 | 1.46 | .196 | Neutral |
|  | difficult | control | double same stimuli | -2.25 | -27.63 | < .001 | Go |
|  |  | inactivation |  | -2.17 | -14.83 | < .001 | Go |
|  | easy | control |  | -0.54 | -10.33 | < .001 | Go |
|  |  | inactivation |  | -0.49 | -7.05 | < .001 | Go |
|  | difficult | control | double different stimuli | 0.01 | 2.64 | .039 | Stay |
|  |  | inactivation |  | 0.02 | 2.93 | .026 | Stay |
|  | easy | control |  | 0.23 | 8.4 | < .001 | Stay |
|  |  | inactivation |  | 0.41 | 5.37 | .002 | Stay |
| M2 | difficult | control | single stimuli | -0.65 | -5.8 | .001 | Go |
|  |  | inactivation |  | -0.38 | -6.5 | .001 | Go |
|  | easy | control |  | 0.15 | 2.25 | .065 | Neutral |
|  |  | inactivation |  | 0.50 | 6.39 | .001 | Stay |
|  | difficult | control | double same stimuli | -1.36 | -19.69 | < .001 | Go |
|  |  | inactivation |  | -1.04 | -9.96 | < .001 | Go |
|  | easy | control |  | -0.18 | -2.14 | .076 | Neutral |
|  |  | inactivation |  | 0.09 | 1.04 | .340 | Neutral |
|  | difficult | control | double different stimuli | 0.08 | 2.79 | .032 | Stay |
|  |  | inactivation |  | 0.27 | 3.91 | .008 | Stay |
|  | easy | control |  | 0.41 | 11.35 | < .001 | Stay |
|  |  | inactivation |  | 0.61 | 9.5 | < .001 | Stay |

**Supplementary Table S8.** Nonparametric tests (Wilcoxon rank sum test) on d-prime and criterion separately for each stimulus type, hemifield and difficulty level. Same as Table 3, but nonparametric. Significant effects are in bold font, consistent effects across the two monkeys are highlighted with gray background.

| Stimulus type | Difficulty | Measure | Hemifield | Monkey 1 |  |  | Monkey 2 |  |  |
| --- | --- | --- | --- | --- | --- | --- | --- | --- | --- |
|  |  |  |  | Wilcoxon rank sum | p-value | Direction of the effect | Wilcoxon rank sum | p-value | Direction of the effect |
| Single stimuli | difficult | criterion | contra | 44 | .318 | - | **34** | **.017** | **Less contra** |
|  |  |  | ipsi | 45 | .383 | - | 53 | .999 | - |
|  |  | d-prime | contra | 59 | .456 | - | 61 | .318 | - |
|  |  |  | ipsi | 57 | .620 | - | 67 | .073 | - |
|  | easy | criterion | contra | 47 | .535 | - | **36** | **.038** | **Less contra** |
|  |  |  | ipsi | 48 | .620 | - | 44 | .318 | - |
|  |  | d-prime | contra | 64 | .165 | - | **70** | **.026** | **Decrease** |
|  |  |  | ipsi | 61 | .318 | - | 60 | .383 | - |
| Double same stimuli | difficult | **criterion** | **contra** | **73** | **.007** | **Less contra** | **74** | **.004** | **Less contra** |
|  |  |  | **ipsi** | **32** | **.007** | **Less contra** | **28** | **.001** | **Less contra** |
|  |  | d-prime | contra | 61 | .318 | - | 63 | .209 | - |
|  |  |  | ipsi | 39 | .097 | - | 44 | .318 | - |
|  | easy | **criterion** | **contra** | **72** | **.011** | **Less contra** | **75** | **.002** | **Less contra** |
|  |  |  | **ipsi** | **32** | **.007** | **Less contra** | **28** | **.001** | **Less contra** |
|  |  | d-prime | contra | 68 | .053 | - | **69** | **.038** | **Decrease** |
|  |  |  | **ipsi** | **32** | **.007** | **Increase** | 52 | .999 | - |
| Double different stimuli | difficult | **criterion** | **contra** | **34** | **.017** | **Less contra** | **74** | **.004** | **Less contra** |
|  |  |  | **ipsi** | **73** | **.007** | Less contra | **52** | .999 | - |
|  |  | d-prime | contra | 58 | .535 | - | **73** | **.007** | **Decrease** |
|  |  |  | ipsi | 62 | .259 | - | **73** | **.007** | **Decrease** |
|  | easy | criterion | contra | 44 | .318 | - | **75** | **.002** | **Less contra** |
|  |  |  | ipsi | 44 | .318 | - | 63 | .209 | - |
|  |  | d-prime | contra | 57 | .259 | - | **75** | **.002** | **Decrease** |
|  |  |  | ipsi | 62 | .620 | - | 68 | .053 | - |

